# Golgi-associated retrograde protein (GARP) complex recruits retromer to trans-Golgi network for FgKex2 and FgSnc1 recycling, necessary for development and pathogenicity of *Fusarium graminearum*

**DOI:** 10.1101/2023.12.07.570498

**Authors:** Yunfei Long, Xin Chen, Jia Chen, Haoran Zhang, Ying Lin, Shuyuan Cheng, Neng Pu, Xuandong Zhou, Renzhi Sheng, Yakubu Saddeeq Abubakar, Huawei Zheng, Yingzi Yun, Guodong Lu, Zonghua Wang, Wenhui Zheng

## Abstract

In eukaryotic cells, the retromer complex plays a crucial role in orchestrating the sorting and retrograde transport of cargo proteins from endosomes to the trans-Golgi network (TGN). Despite its significance, the molecular details of this intracellular trafficking process remain unclear. Here, we identified a Golgi-associated retrograde protein (GARP) complex as a mediator of vesicular transport, facilitating the recruitment of the retromer complex to TGN to exert its functions. The GARP complex is mainly localized to the TGN, where it interacts with the retromer complex. This interaction is evolutionarily conserved across different species. Furthermore, we identified FgKex2 and Fgsnc1 as cargo proteins in the GARP/retromer-mediated recycling pathway. Loss of GARP or retromer results in a complete mis-sorting of FgKex2 and FgSnc1 to the vacuolar degradation pathway, which affects the growth, development, toxisome biogenesis, and pathogenicity of *F. graminearum*. In summary, we assert that GARP facilitates the recruitment of retromer from endosomes to the TGN, orchestrating the recycling of FgKex2 and FgSnc1. This is integral to sustaining continuous growth, development and contributes significantly to the pathogenicity of *F. graminearum*.

## Introduction

*Fusarium graminearum* ranks among the top 10 fungal pathogens affecting plants, causing Fusarium head blight (FHB). This disease poses a significant threat to both the quality and yield of wheat and other small grain cereal crops on a global scale^1–3^. The fungus invades its hosts by exploiting vulnerable openings or through stomata^4^. Furthermore, this fungus is recognized for its ability to form lobate appressoria and infection cushions as part of the infection processes^5,6^. Following this, it initiates colonization within host cells through hyphal elongation and apical growth^7^. Grains from infected cereals are typically unsuitable for consumption, as the pathogen releases perilous mycotoxins like zearalenone and deoxynivalenol (DON), which result in grain contamination^8^. Today, FHB has become a worldwide cause for concern, inflicting substantial economic losses and jeopardizing global food security^9^. This underscores the pressing necessity to address this issue. A comprehensive grasp of the biology and molecular mechanisms governing *F. graminearum* pathogenesis will provide valuable insights into potential drug targets that hold promise for effectively controlling and managing FHB.

In eukaryotic cells, vesicle transport stands as a pivotal process orchestrating numerous cellular functions^10^. The exchange of macromolecules (including proteins and lipids) among the subcellular organelles primarily relies on vesicle transport ^11^. This intricate journey encompasses key stages such as budding, transport, tethering, anchoring, and membrane fusion^12^. The tethering factor plays a pivotal role in initiating contact between the transport vesicles and the receptor membrane^10,13^. It identifies the transport vesicles and brings them to close contact with the receptor membrane^14^. During this process, the tethering factor, in conjunction with small GTPases and their downstream partners, interacts with some SNARE proteins to collectively regulate the membrane fusion process^15,16^. Presently, there are a total of ten multi-subunit tethered complexes, strategically positioned within different transport pathways, each carrying out its distinct regulatory functions^17,18^. Among these, the Golgi-associated retrograde protein (GARP) complex primarily resides within the trans-Golgi network (TGN), functioning as a tethering factor that mediates retrograde transport from endosomes to the TGN^14,19^.

The GARP complex was first identified in the context of yeast-based investigations into protein transport mechanisms^15,20^. Classified among the multi-subunit tethering complexes (MTCs), GARP is positioned at the periphery of the cytosolic face of the TGN. As a heteromeric protein complex, it consists of four distinct subunits: vacuolar protein sorting 51 (VPS51), VPS52, VPS53 and VPS54^21^. The GARP complex plays a crucial role in the tethering transport vesicles with the TGN, ultimately facilitating the fusion between the vesicles and the receptor membranes^14^. Fusion of transport vesicles with the TGN is instrumental in promoting the recycling of various membrane proteins, such as the vacuolar protein sorting receptor Vps10^22^, the cell wall sensor Wsc1^23^. Similar to yeast, the mammalian GARP complex predominantly localizes within the TGN and is required for the retrieval of lysosomal hydrolase precursor receptors, such as mannose 6-phosphate receptors (MPRs), from endosomes to the TGN^24,25^. In Arabidopsis, the homologous genes encoding the Vps52/53/54 subunits are *POK*/*AtVPS52*, *HIT1*/*AtVPS53* and *AtVPS54*^26,27^. Unh protein, a homologue of Vps51, interacts with Vps52 and is localized to the TGN and pre-vacuole regions. Mutation of the *UNH* gene results in weakened directional transport of vacuoles, leading to altered leaf morphology and reduced root and shoot growths in the plant^28^. Information regarding GARP in filamentous fungi is scarce; a singular report in Botrytis cinerea reveals that exogenous small RNAs (sRNAs) can suppress the expression of VPS51, consequently influencing the virulence of B. cinerea^29^. While the GARP complex is remarkably conserved across species, its functional intricacies have remained unexplored especially in pathogenic fungi.

In addition to the GARP, some transport machinery, such as the retromer complex, is needed to transport cargo proteins back to the right location for their biological functions^10,30^. The retromer protein complex mediates the vesicular transport of some membrane proteins from the endosomes to the TGN and the plasma membrane^31^. It is a heteropentamer constructed from two major subcomplexes. The core subcomplex is a trimer composed of Vps35, Vps29 and Vps26, commonly known as cargo selection complex (CSC); another component is the dimer of sorting nexin (SNX) proteins (such as Vps5 and Vps17), which binds to phosphatidylinositol 3-phosphate-enriched endosomal membranes during cargo sorting^9^. Recently, we reported that in plant pathogenic fungi, including rice blast fungus (*Magnaporthe oryzae*) and *F. graminearum*, retromer complex is involved in endosomal sorting, regulating developmental and pathogenic processes by collecting and recycling specific cargo proteins^32,33^. Nevertheless, the association between the GARP complex and the retromer complex remains entirely unexplored in eukaryotes, including model organisms such as yeast and mammals.

In this study, we found that the GARP complex components localized to TGN and loss of GARP significantly impairs growth, conidiation, virulence, and impeded DON production of *F. graminearum*, highlighting its crucial role in fugal development and pathogenicity. We further identified the retromer complex as an interacting partner of GARP and the interaction model between GAPR and retromer was established. Remarkably, deletion of GARP disrupted the TGN localization of retromer, suggesting the important role of GARP in recruitment of retromer to TGN structure. This further ensures the proper function of GARP/retromer-mediated vesicle trafficking process. We further identified FgKex2 and FgSnc1 as cargo proteins of GARP/retromer-mediated retrograded machinery. Mis-organization of the FgKex2 and FgSnc1, caused by the disruption of GARP/retromer trafficking machinery, further impaired the development and pathogenicity of *F. graminearum*. In summary, we presented the first systematic functional study of the GARP complex in a phytopathogenic fungus and unveiling its interaction with the retromer complex.

## Materials and Methods

### Strains and culture conditions

In this study, the *Fusarium graminearum* wild-type (WT) strain PH-1 was used as a control, and also served as the background from which all the mutant strains were generated (see supporting information Table S1. All incubations of both PH-1 and the mutant strains were carried out at a temperature of 28°C for 3 days on either solid or liquid complete medium (CM), unless otherwise stated. Determination of growth rates on CM plates and conidiation in liquid carboxymethylcellulose (CMC) medium were achieved following previously established protocols^33^. Assessment of sexual reproduction was also performed in accordance with a methodology outlined in a previous report ^34^.

### Fungal transformation and generation of gene deletion mutants

Protoplast preparation and fungal transformation of *F. graminearum* were conducted following standard procedures ^35^. Gene deletion mutants were generated for all GARP complex subunits and *FgKEX2* genes by split-marker approach ^36^. The primers used to amplify the flanking sequences for each gene are listed in Table S2 and the resulting transformants were screened by PCR and further confirmed by Southern blot.

### Vector construction

Unless specified otherwise, all vectors were constructed by amplifying an entire protein coding sequences, including the native promoter, using the primers listed in Table S3. These sequences were then fused with green fluorescent protein (GFP), mCherry Fluorescent Protein (mCherry), C-terminal yellow fluorescent protein (CYFP), or N-terminal yellow fluorescent protein (NYFP), as the case may be, using One-Step Cloning Kit (C112-01, Vazyme, Nanjing, China), following the manufacturer’s protocol. Details of the plasmid construction approach in this study are presented in (Table S3).

### Pathogenicity and DON Production Assays

As previously outlined^37^, we assessed the fungal virulence on wheat spikelets, coleoptiles and leaves. Symptoms on the infected flowering wheat heads were observed at 14 days post-infection (dpi). Additionally, symptoms on wheat coleoptiles and leaves were evaluated at 7 dpi. The amount of deoxynivalenol (DON) produced by each strain was quantified using a DON detection kit (Wiseste Biotech Co. Ltd, China) after a week of incubation in liquid trichothecene biosynthesis induction (TBI) medium. Quantitative real-time PCR (qRT-PCR) was conducted using ChamQ SYBR qPCR Master Mix (Q311-02, Vazyme, China) to determine the mRNA levels of TRI genes, and the primer details for the qRT-PCR assay are provided in (Table S3).

### Yeast two-hybrid (Y2H) and bimolecular fluorescence complementation (BiFC) assays

The Y2H assay was conducted following established protocols ^33^. In summary, full-length cDNAs of the target genes were inserted into pGADT7 (AD) containing GAL4 activation domain and pGBKT7 (BD) containing GAL4-binding domain to create bait and prey constructs, respectively. Subsequently, the constructs were transformed into *Saccharomyces cerevisiae* strain AH109 strain, and the transformants were grown on synthetic medium (SD) containing-Trp-Leu-His-Ade. Interaction of BD-Lam with AD-T was used as negative control, while that of BD-53 with AD-T served as positive control. For BiFC assays, *FgVPS51*, *FgVPS53* and *FgVPS35* gene fragments were cloned into a pKNT-NYFP (N-terminal portion of the YFP fluorescent protein) vector, while *FgKEX2* and *FgVPS35* fragments were cloned into a pKNT-CYFP (C-terminal portion of the YFP fluorescent protein) vector. Strains expressing NYFP and/or CYFP, including FgVps51-NYFP, FgVps53-NYFP, FgVps35-NYFP, FgVps51-NYFP+CYFP, FgVps53-NYFP+CYFP, FgVps35-NYFP+CYFP, FgVps35-CYFP, FgKex2-CYFP, FgVps35-CYFP+NYFP, FgKex2-CYFP+NYFP, were used as negative controls. Hygromycin- and/or neomycin-resistant transformants were isolated and confirmed by PCR. YFP signals were visualized using a Nikon A1 laser confocal microscope (Nikon, Japan).

### Sexual reproduction assays

To induce sexual reproduction, strains were cultured on carrot agar medium (carrot 200 g/L, agar 16 g/L) for a period of 3 days. The mycelia that developed on the carrot agar surface were gently scraped off, and 1 ml of 2.5% Tween 20 solution was added. The cultures were further incubated at 25°C for 10 days based on a previously established procedure^38^. Perithecia formation and ascospores were observed two weeks afterwards. Each experiment was independently repeated three times.

### Affinity capture-mass spectrometry analysis

Conidia were harvested from the strains expressing FgVps51-GFP, FgVps53-GFP, FgVps35-GFP, and PH-1-GFP. Subsequently, they were suspended in 500 ml of liquid CM medium at a concentration of 10 × 10^4^ conidia per milliliter and incubated at 28 °C for 16 hours, in a rotary shaker operating at 110 rpm. The mycelia were collected, washed with sterile ddH_2_O, ground into fine powder in liquid nitrogen, and suspended in 10 ml of extraction buffer containing protease inhibitors (10 mM Tris/Cl pH 7.5, 150 mM NaCl, 0.5 mM EDTA, 0.5% NP40), followed by incubation at 4°C for 30 minutes. The lysate was then centrifuged at 20,000 g at 4 ℃ for 20 minutes. The resulting supernatant was transferred to GFP-Trap A beads at a ratio of 10 ml to 25 μl beads and gently rotated at 4℃ for approximately 3 hours. Subsequently, affinity purification experiments were conducted following the procedures outlined in a previous report^37^.

### Staining and live cell imaging of *F. graminearum*

FM4-64 (Molecular Probes, Eugene, OR, USA, final concentration 8 μM) was used to stain the fungal hyphae to visualize the Spitzenkorper, plasma membrane, septum, and vacuolar membrane. Fresh conidia were stained with 0.1 mg/ml calcofluor white (CFW) (Sigma-Aldrich, USA) for 30 seconds to visualize septa. For subcellular localization assays using mycelia, a mycelial block from CM or SYM agar containing the leading hyphae was excised and placed upside down on a coverslip and observed directly under a confocal microscope. The excitation wavelengths used were 488 nm for GFP, 561 nm for mCherry, and FM4-64, and 405 nm for CFW. Live cell fluorescence imaging was conducted using a Nikon A1R laser scanning confocal microscope system (Nikon, Japan). Sequential firing of lasers for the channels to be used was performed to prevent cross-excitation, as previously described^33^. All microscope images were captured within a single focal plane unless noted otherwise, and sequence images were exported as AVI files.

### Protein structure prediction and molecular docking

The protein sequences of FgKex2, FgSnc1, FgVps53, FgVps35, MoVps53, MoVps35, FoVps53, and FoVps35 were obtained from NCBI database (https://www.ncbi.nlm.nih.gov/). The protein structures of FgKex2, FgSnc1, FgVps53, FgVps35, MoVps53, MoVps35, FoVps53, FoVps35 ScVps53, and ScVps35were predicted using A_LPHA_F_OLD2_^39^. Molecular docking analyses of FgVps53 with FgVps35, MoVps53 with MoVps35, and FoVps53 with FoVps35 were conducted using GRAMM (https://gramm.compbio.ku.edu/)^40^.

## Results

### Identification and deletion of *FgVPS51* gene in *F. graminearum*

The gene coding the core subunit of the GARP complex, *FgVPS51* (FGSG_06320), was identified through BLASTp analysis at the *F. graminearum* genome database (https://fungidb.org/fungidb/), using the *S. cerevisiae* Vps51 protein sequence as a query. The *FgVPS51* gene spans 932 base pairs, encoding a 292-amino acid protein. The similarity of FgVps51 protein to its orthologues in fungi, plants and humans is relatively high, ranging from 93% in *Fusarium oxysporum* (FoVps51) to 45% in *Arabidopsis thaliana* (Vps51) (Fig. S1). Phylogenetic analysis further supports the ancient origin and exceptional conservation of the Vps51 protein across various species (Fig. S1). To analyze the biological functions of FgVps51 in *F. graminearum*, we generated *FgVPS51* gene deletion mutants by replacing the *FgVPS51* coding sequence with a hygromycin phosphotransferase (hph) gene as a selective marker. Southern blot analysis confirmed the mutants, revealing a 4.62-kb band in the wild-type PH-1 and a 2.20-kb band in the Δ*Fgvps51* mutant (Fig. S2). Additionally, a FgVps51-GFP vector was constructed (under the *FgVPS51* native promoter) and transformed into the Δ*Fgvps51* mutant, resulting in the generation of a complemented strain (Δ*Fgvps51-C*).

### FgVps51 predominantly localizes to the TGN

A previous research has established that Vps51 predominantly localizes to the TGN in yeast ^14^. In our investigation of the localization of FgVps51-GFP in *F. graminearum*, we found that the FgVps51-GFP primarily exhibited a punctate cytoplasmic distribution at various developmental stages, ranging from conidiogenesis (from phialides to mature conidia) to germination (from germlings to vegetative hyphae). Notably, the FgVps51-GFP signal was strongly expressed in vegetative hyphae and matured conidia (Fig. S3). To further investigate the subcellular localization of FgVps51 in *F. graminearum*, we generated constructs harboring the early endosomal marker mCherry-FgRab52^34^ and the TGN marker FgSec7-mCherry^41,42^. These constructs were co-transformed with FgVps51-GFP into the Δ*Fgvps51* deletion mutant, respectively, and their intracellular localizations were examined by confocal microscopy. As illustrated in (Fig. 1a, b), we observed that FgVps51-GFP co-localized significantly with FgSec7-mCherry (65.58±7.06% colocalization) and partially with mCherry-FgRab52 (12.43±2.79% colocalization) in hyphal cells. To assess whether FgVps51-GFP co-localized with the vacuole membrane, we used the fluorescent dye FM4-64 to mark membrane internalization and transport to the vacuolar membranes. Our data indicated that FgVps51-GFP partially colocalizes with the vacuolar membrane-selective dye FM4-64 (13.88±3.86% colocalization) in hyphal cells (Fig. 1c). As such, we conclude that FgVps51 predominantly localizes to the TGN.

**Fig. 1.**
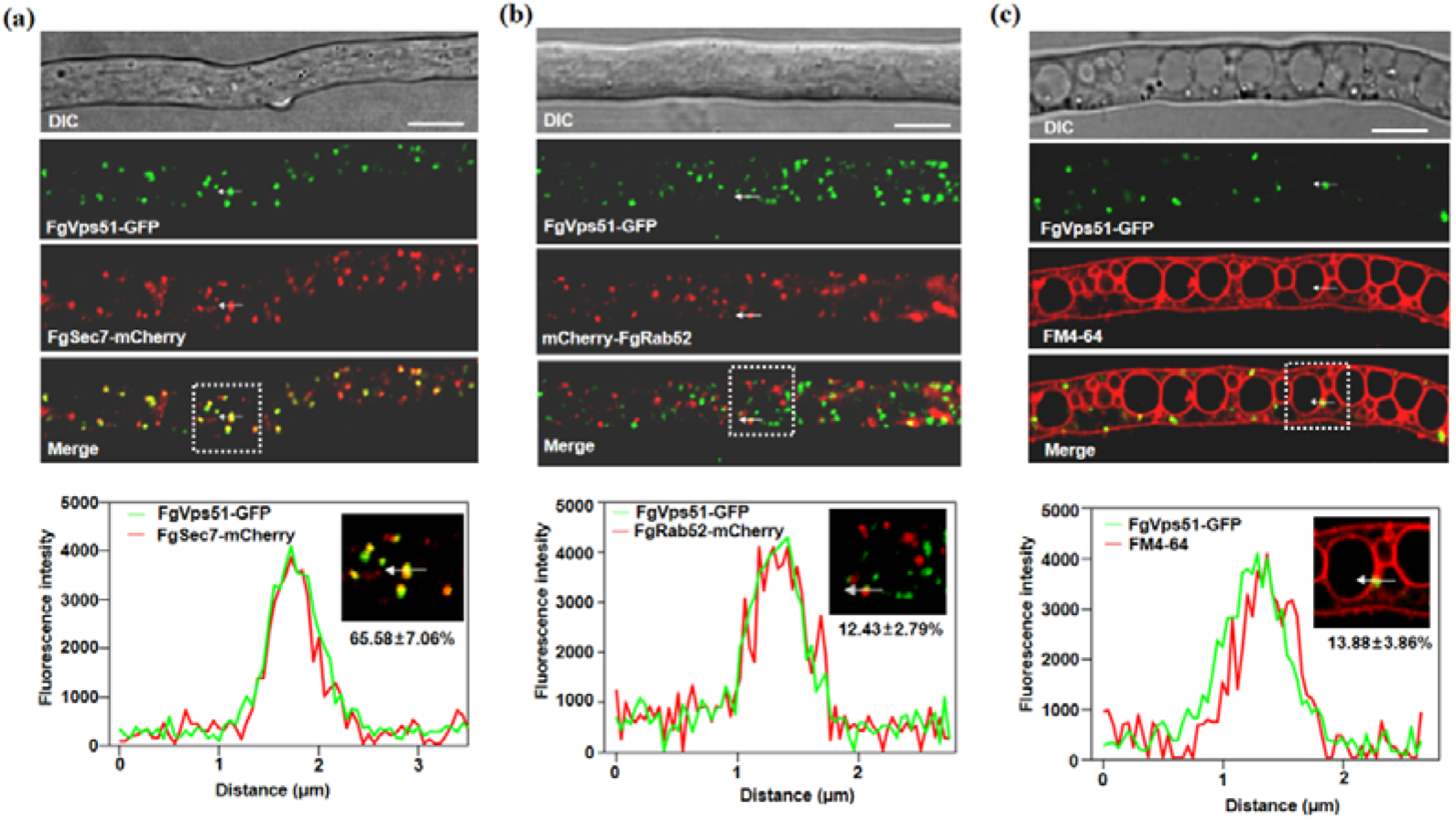
FgVps51-GFP mainly localizes to trans-Golgi network in *Fusarium graminearum*. (a) FgVps51-GFP largely co-localized with the trans-Golgi network (TGN) marker FgSec7-mCherry. The co-localization rate is 65.58 ± 7.06%. (b) FgVps51-GFP partially co-localized with the early endosome marker mCherry-FgRab52. The co-localization rate is 12.43 ± 2.79%. (c) FgVps51-GFP partially co-localized with the vacuole membrane dye FM4-64. The co-localization rate is 13.88 ± 3.86%. Each line scan graph was generated at the position indicated by the arrow to show the relative co-localization of FgVps51-GFP (green) with FgSec7-mCherry (red), mCherry-FgRab52 (red) and FM4-64 (red), respectively. Bar=10 µm. DIC, differential interference contrast.

### The GARP complex subunit FgVps51 is essential for hyphal growth, conidiation and sexual development

To examine the biological functions of FgVps51 in *F. graminearum*, the wild-type strain PH-1, the *FgVPS51* gene deletion mutant (Δ*Fgvps51*) and the complemented strain (Δ*Fgvps51-C*) were cultured on CM at 25°C for 3 days. Colony morphology and diameters of each strain were measured and analyzed. The results revealed that the Δ*Fgvps51* mutant exhibited significantly slower growth compared to the wild-type strain PH-1 and the complemented strain Δ*Fgvps51-C* (Fig. 2a, b). Notably, lush and bushy aerial hyphae were observed in the colonies of the wild-type and the complemented strains, while the Δ*Fgvps51* mutant exhibited rare and barren aerial hyphae (Fig. 2c, d). Furthermore, we closely observed the growing hyphae of the various strains using a confocal microscope. As depicted in (Fig. 2e), hyphae from the Δ*Fgvps51* mutant strain failed to maintain stable polarized growth, displaying curved growth compared to the straight-growing hyphae of the wild-type and the complemented strain. Therefore, we conclude that FgVps51 is essential for the normal vegetative growth of *F. graminearum*.

**Fig. 2.**
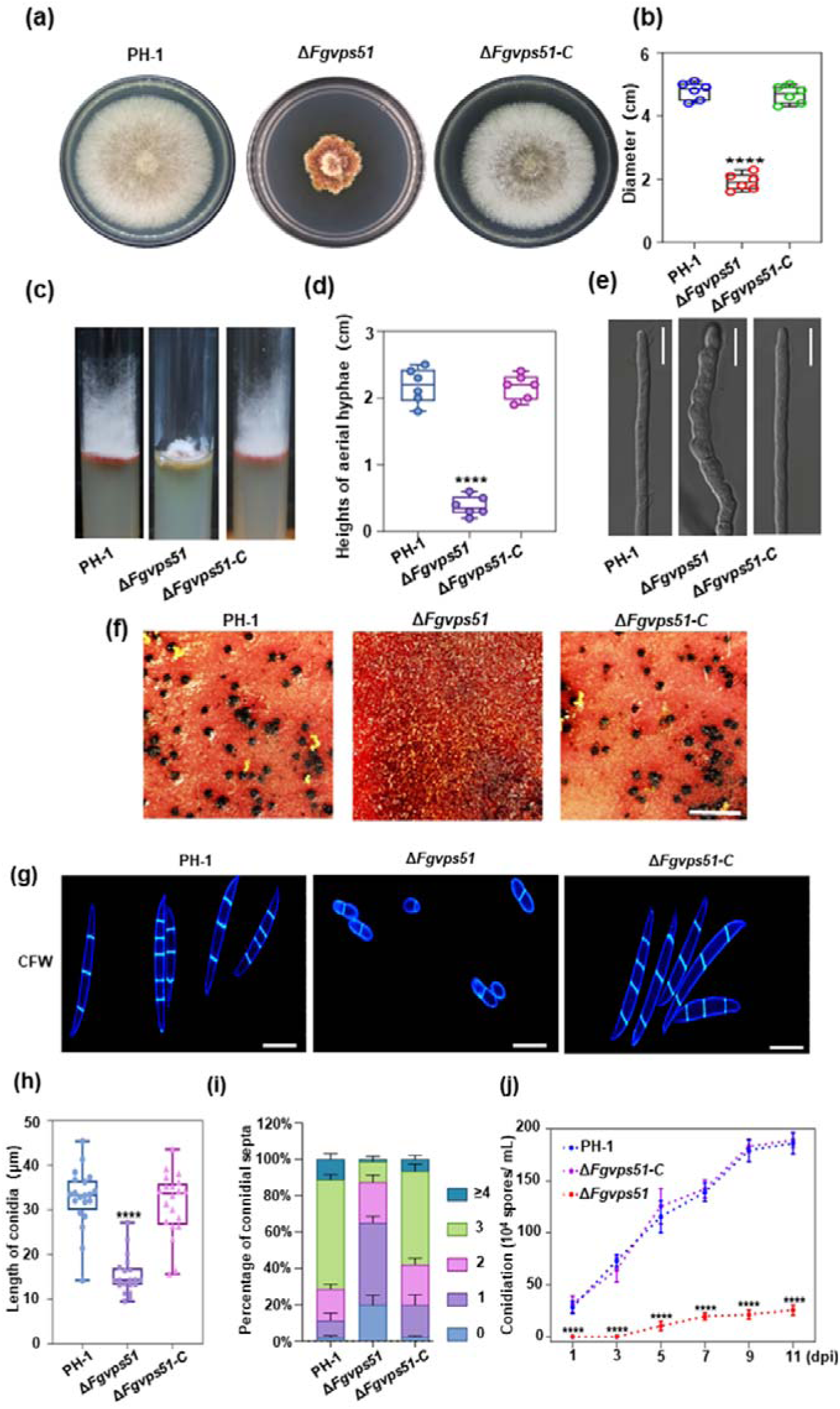
FgVps51 is essential for the vegetative growth and asexual development of *F. graminearum.* (a) The wild type (PH-1), *FgVPS51* deletion mutant (Δ*Fgvps51*) and complemented strain (Δ*Fgvps51-C*) were cultured on complete medium (CM) at 28 ℃ for 3 days. (b) The colony diameters of PH-1, Δ*Fgvps51* and Δ*Fgvps51-C* after culture on CM for 3 days. (c) Aerial hyphal growths of the indicated strains cultured in glass tubes containing CM. (d) The heights of the aerial hyphae from the tested strains in cylindrical glass tubes. (e) Microscopic morphology of the hyphae from the indicated strains. Bar=10 µm. (f) Conidial morphology and septation for PH-1, Δ*Fgvps51* and Δ*Fgvps51-C* strains after staining with 10 µg/mL calcofluor white (CFW). Bar=10 µm. (g) The lengths of the conidia from the tested strains. (h) Statistical analysis of the number of septa in the conidia produced by the indicated strains. (i) The PH-1, Δ*Fgvps51* and Δ*Fgvps51-C* conidia harvested from liquid carboxymethylcellulose (CMC) medium at different time points after inoculation. (j) Perithecial formation by the indicated strains after growth on carrot agar media for 2 weeks. Bar=1 mm. The values shown are means of independent experiments. Statistical analysis was processed by two-way ANOVA for multiple comparisons using GraphPad Prism 9 (****p < 0.0001).

Asexual conidia and sexual ascospores produced by *F. graminearum* are recognized as crucial inocula for infecting flowering wheat heads ^43,44^. Consequently, we investigated the roles of FgVps51 in conidiogenesis and sexual reproduction of *F. graminearum*. Initially, the wild-type PH-1, the mutant Δ*Fgvps51*, and the complemented Δ*FgVps51-C* strains were incubated in liquid carboxymethylcellulose (CMC) media for conidia production. After 5 days of incubation, we obtained 115.4 ± 15.53 × 10^4^ conidia ml^−1^ and 125.6 ± 16.20 × 10^4^ conidia ml^−1^ from the PH-1 and complemented Δ*FgVps51-C* strains, contrary to 10.40 ± 4.72 × 10^4^ conidia ml^−1^ obtained from the Δ*Fgvps51* strain. Interestingly, conidia production by the PH-1 and the complemented strains increased with increase in incubation time, reaching their maxima of 184.60 ± 10.53 × 10^4^ conidia ml^−1^ and 189.40 ± 7.00 × 10^4^ conidia ml^−1^, respectively, at the 11^th^ day. Conversely, the conidia production of the Δ*Fgvps51* mutant did not significantly change with increase in incubation time (Fig. 2j). We further stained the conidia with CFW (10 μg/mL) to visualize their septa under a fluorescence microscope. The mutants’ conidia had fewer number of septa than those of the PH-1 and complemented strains, as shown in (Fig. 2g, i). The average length of the conidia produced by the Δ*Fgvps51* mutant was only 11.97 ± 1.68 μm, while those from the PH-1 and complemented strain had average lengths of 35.14 ± 2.79 μm and 34.84 ± 2.41 μm, respectively (Fig. 2h). Additionally, the wild-type strain PH-1 produced a large number of perithecia after 2 weeks of culture on carrot medium, while no perithecia were observed for Δ*Fgvps51* mutant under the same conditions (Fig. 3f). These data indicate that FgVps51 is crucial for normal vegetative growth, conidiogenesis, conidial morphology and sexual reproduction in *F. graminearum*.

**Fig. 3.**
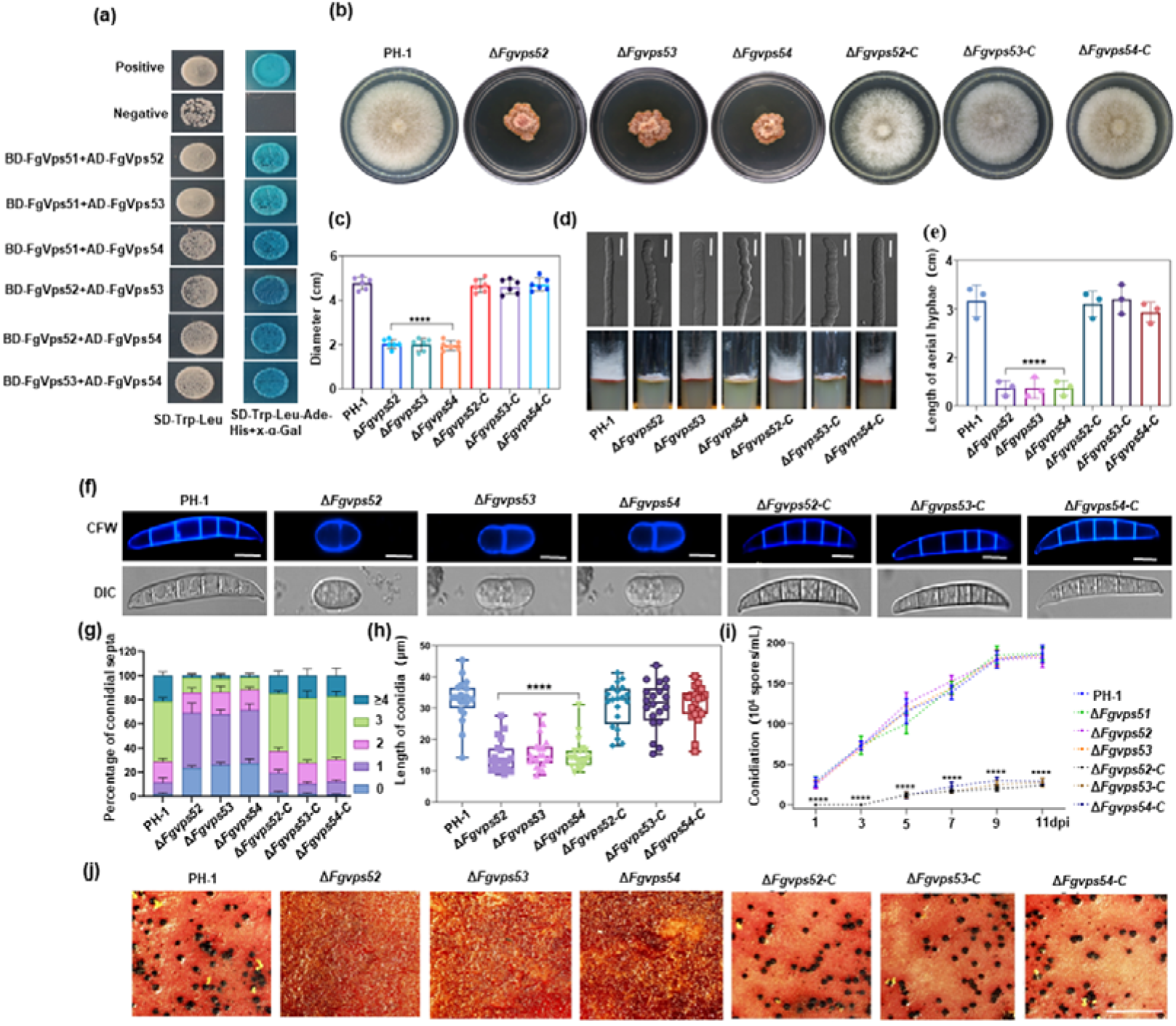
Components of GARP complex are necessary for hyphal growth, conidial development and ascospore formation in *F. graminearum.* (a) Yeast two hybrid (Y2H) assay showing positive interactions among the GARP components (FgVps51, FgVps52, FgVps53 and FgVps54) in *F. graminearum*. (b) Colonies of the wild type (PH-1), each gene deletion mutant (Δ*Fgvps52,* Δ*Fgvps53 and* Δ*Fgvps54*), and their respective complemented strains (Δ*Fgvps52-C,* Δ*Fgvps53-C and* Δ*Fgvps54-C*) grown on CM media for 3 d. (C) Colony diameters of the tested strains grown on CM media under same experimental conditions. (d) Aerial hyphae of the indicated strains grown on CM in cylindrical glass tubes. Bar=10 µm. (e) Statistical analyses of the heights of aerial hyphae of the tested strains grown on the CM in cylindrical glass tubes. (f) Conidia of the indicated strains stained with CFW (calcofluor white) to visualize septa. Bar=10 µm. (g) Percentage of conidia with different number of septa in the tested strains. (h) Statistical analysis of the lengths of conidia from the tested strains. (i) The number of conidia produced by the GARP component mutants is significantly reduced at different time points. (j) Perithecia production ability of the indicated strains after 2 weeks of inoculation on carrot agar plates. Bar=1 mm. The values shown are means of independent experiments. Statistical analysis was processed by two-way ANOVA for multiple comparisons using GraphPad Prism 9 (****p < 0.0001).

### Other subunits of the GARP complex are conserved, demonstrating mutual interaction in *F. graminearum*

Apart from Vps51, the yeast GARP complex comprises three additional vacuolar protein sorting proteins namely Vps52, Vps53 and Vps54 which also localize primarily in the TGN^14,20^. To identify the homologues of these three GARP complex subunits in *F. graminearum* genome, we carried out BLASTp analyses of their respective protein sequences in yeast against the *F. graminearum* genome. These analyses revealed the homologues of yeast *VPS52*, *VPS53* and *VPS54* as FGSG_01954 (*FgVPS52*), FGSG_00870 (*FgVPS53*) and FGSG_06778 (*FgVPS54*), respectively, in *F. graminearum*. Phylogenetic analyses of FgVps52, FgVps53 and FgVps54 across fungi, plants and mammals indicated their ancient origin and exceptional conservation (Fig. S4, S5, S6). Current studies propose that the GARP complex’s four subunits assemble through a four-helix bundle, forming a multisubunit tethering complex (MTC)^13^. However, the interaction pattern among the GARP subunits in plant pathogenic fungi remains unclear. To investigate these interactions in *F. graminearum*, each of the four GARP components was respectively cloned into pGADT7 (AD) vector containing a Gal4 activation domain or pGBKT7 (BD) vector containing a DNA-binding domain, and a yeast two-hybrid (Y2H) assay was conducted. As shown in (Fig. 3a), each individual subunit of the GARP complex showed positive interaction with the other three subunits.

### The GARP subunits FgVps52, FgVps53 and FgVps54 are required for hyphal growth, conidiation and Sexual development

To elucidate the functions of the three remaining GARP complex subunits in *F. graminearum*, we employed a targeted gene replacement strategy to delete *FgVPS52*, *FgVPS53* and *FgVPS54* genes in the wild-type PH-1. All gene deletion mutants were confirmed through PCR and subjected to Southern blots (Fig. S7). Additionally, we generated gene complementation strains by reintroducing the open reading frame (ORF) of the three GARP complex genes, each with its native promoter fused with green fluorescent protein (GFP) into the respective deletion mutants, and successfully generated all complemented strains. To further validate the essential role of the GARP subunits FgVps52, FgVps53 and FgVps54 in hyphal growth, conidiation and sexual development, all the strains were inoculated on CM and incubated at 28°C. As shown in (Fig. 3b, c, d, e), after 3 days of incubation, all the complemented strains had phenotypes similar to the wild-type. However, like the Δ*Fgvps51* strain, the mutant strains Δ*Fgvps52*, Δ*Fgvps53* and Δ*Fgvps54* exhibited noticeable defects in colony morphology, diameters, aerial hyphae, and mycelial microstructure. Furthermore, we investigated the roles of the GARP complex components FgVps52, FgVps53 and FgVps54 in asexual and sexual developments. Our results demonstrated that the complemented strains and the wild-type strain PH-1 produced normal sickle-shaped conidia. However, the absence of the GARP complex subunits resulted in reduced number of septa in *F. graminearum*, as evidenced by CFW staining of their respective conidia (Fig. 3f, g). Additionally, the lengths of the conidia in the mutant strains were significantly reduced compared to the PH-1 and the complementation strains (Fig. 3h). Intriguingly, when all the strains were cultured in carboxymethyl cellulose (CMC) liquid media, the number of conidia produced by the mutant strains was significantly lower than that of PH-1 and complementation strains at different incubation time points (Fig. 3i). To confirm the role of the GARP complex in sexual development, the wild-type PH-1, the gene deletion mutants (Δ*Fgvps52*, Δ*Fgvps53* and Δ*Fgvps54*), and all the complemented strains were inoculated on carrot media for 2 weeks. As shown in (Fig. 3j), no perithecia were observed in the GARP complex mutants under the same conditions. Taken together, these results lead us to concluding that the GARP complex is crucial for hyphal growth, conidiation and sexual development in *F. graminearum*.

### GARP complex primarily localizes to the TGN

To ascertain the localization of the GARP complex, we analyzed the subcellular localizations of FgVps52-GFP, FgVps53-GFP and FgVps54-GFP subunits during conidiogenesis, germina conidia, in matured conidia and in growing hyphae by confocal microscopy. Our observations demonstrated that FgVps52-GFP, FgVps53-GFP and FgVps54-GFP displayed punctate distribution at the various developmental stages and under different culture conditions (Fig. S8). Since our earlier data showed that the core GARP subunit FgVps51-GFP primarily localizes in the TGN, we further investigated whether the other subunits share a similar localization. To achieve this, we co-transformed the Golgi marker FgSec7-mCherry with FgVps52-GFP, FgVps53-GFP and FgVps54-GFP, respectively, into the GARP complex mutants. Confocal microscopy revealed that each of FgVps52-GFP, FgVps53-GFP and FgVps54-GFP co-localized with the TGN marker (Fig. 4a, b, c). Hence, these findings provide conclusive evidence that the GARP complex localizes to the TGN in *F. graminearum*.

**Fig. 4.**
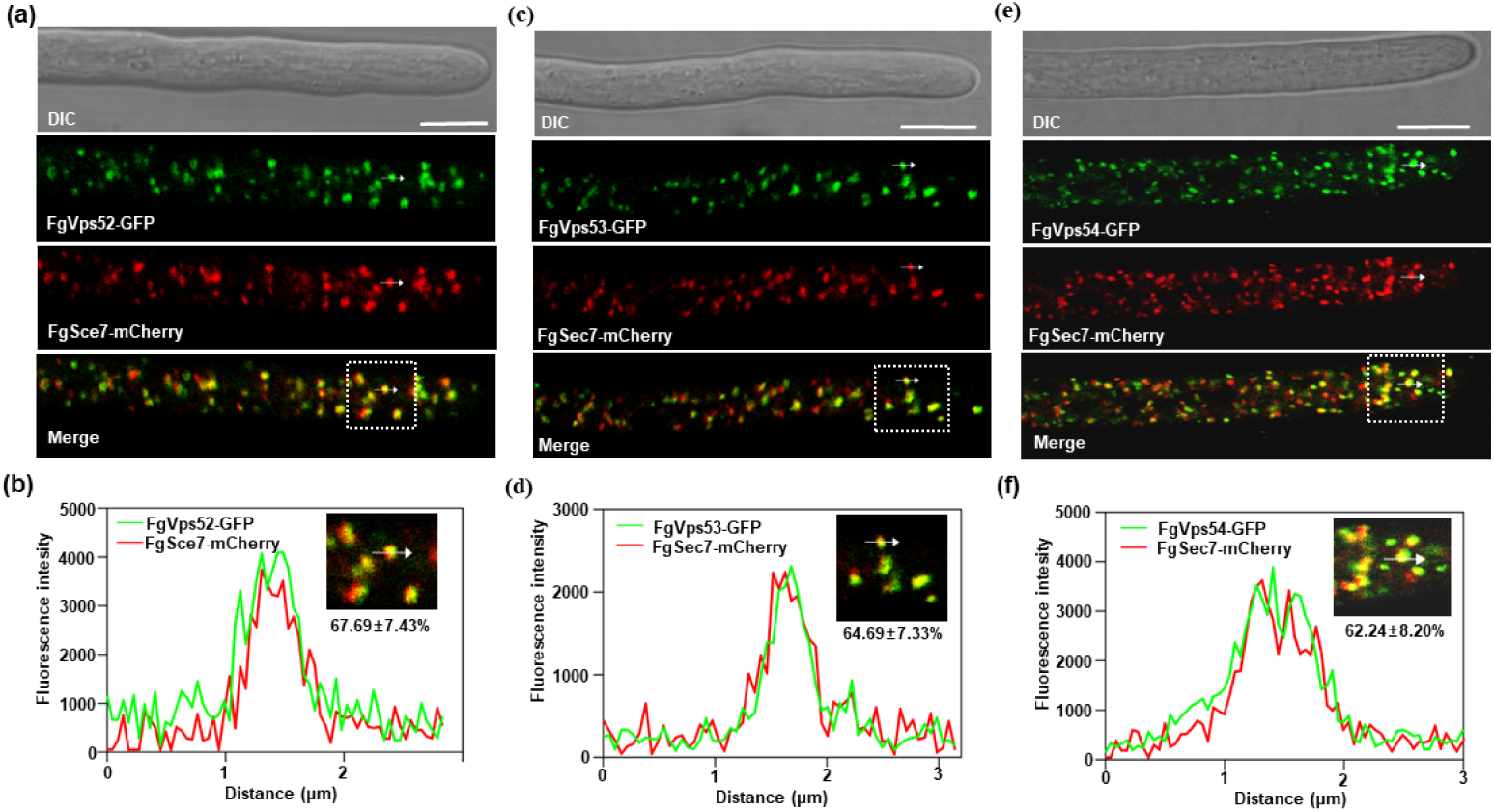
Co-localization of GARP components with the TGN in *F. graminearum*. (a) FgVps52 co-localized with FgSec7-positive TGN (67.69 ± 7.43% co-localization). (b) FgVps53 co-localized with FgSec7-positive TGN (64.69 ± 7.33% co-localization). (c) FgVps54 co-localized with FgSec7-positive TGN (62.24 ± 8.20% co-localization). Each line scan graph was generated at the position indicated by the arrows to show the relative co-localization of FgVps52-GFP (green), FgVps53-GFP (green) and FgVps54-GFP (green) with FgSec7-mCherry (red), respectively. Bar=10 µm. DIC, differential interference contrast.

### GARP complex is indispensable for pathogenicity and deoxynivalenol (DON) production

To explore the role of the GARP complex in the pathogenesis of *F. graminearum*, we initially assessed the virulence of the GARP complex mutants in flowering wheat heads. The results indicated that the mutants Δ*Fgvps51*, Δ*Fgvps52*, Δ*Fgvps53* and Δ*Fgvps54* of the GARP complex caused fewer blight symptoms on spikelets compared to PH-1 and complemented strains (Fig. 5a). Furthermore, when the strains were inoculated on wheat coleoptiles and leaves, compared to PH-1 and complemented strains, the GARP complex mutants caused significantly reduced disease symptoms after 7 days of incubation under moist conditions (Fig. 5a, S9). Moreover, the GARP complex mutants displayed markedly lower disease index scores than PH-1 and complemented strains (Table S4). These findings collectively demonstrate that the GARP complex contributes significantly to the virulence of *F. graminearum*.

**Fig. 5.**
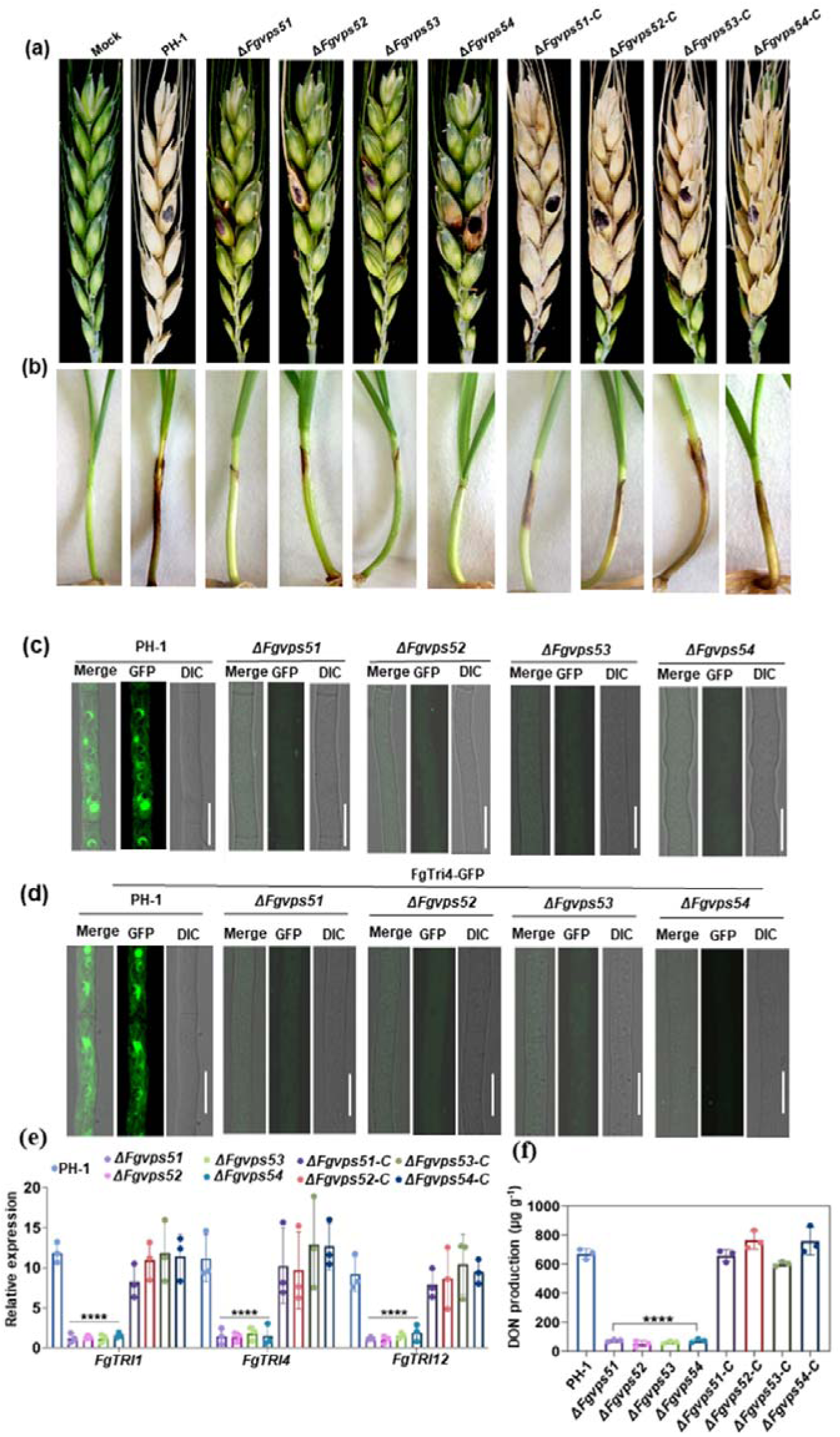
The GARP complex is important for *F. graminearum* virulence. (a) Flowering wheat heads were inoculated with mycelial plugs from the PH-1, GARP component mutants and their respective complemented strains. The number of diseased spikelets per wheat head was measured and photographed at 14 dpi (days post infection). (b) The pathogenicity assay of the tested strains on wheat coleoptiles and the lesion lengths were examined at 7 dpi. (c) Localization of the DON biosynthesis protein FgTri1-GFP in PH-1 and GARP component mutants. Bar = 10 μm. (d) Localization of the DON biosynthesis protein FgTri4-GFP in PH-1 and GARP component mutants. Bar = 10 μm. (e) Bar charts showing relative expressions of *FgTRI1*, *FgTRI4* and *FgTRI12* genes in the tested strains. (f) Bar charts showing DON production potential of the indicated strains in liquid trichothecene biosynthesis induction (TBI) media. The values shown are means of independent experiments. Statistical analysis was processed by two-way ANOVA for multiple comparisons using GraphPad Prism 9 (LLLLp < 0.0001).

As a mycotoxin and virulence factor, deoxynivalenol (DON) plays a crucial role in the infection of *F. graminearum*^45^. To determine whether the GARP complex is necessary for DON production, the PH-1, all gene deletion mutants and complemented strains were cultured in liquid trichothecene biosynthesis induction (TBI) media at 28°C for 7 days in the dark. As shown in (Fig. 5f), production of DON in the mutants Δ*Fgvps51*, Δ*Fgvps52*, Δ*Fgvps53* and Δ*Fgvps54* was significantly reduced compared to PH-1 and complemented strains. Subsequently, we assessed the expression levels of some DON biosynthesis genes in the mutant strains cultured in TBI media using quantitative real-time polymerase chain reaction (qRT-PCR). The results indicated that the expression levels of three genes *FgTRI1, FgTRI4* and *FgTRI12* related to the biosynthesis of trichothecenes were significantly down-regulated in the GARP component mutants compared to PH-1 and complemented strains (Fig. 5e). To further corroborate our results, since FgTri1 and FgTri4 are toxisome marker proteins ^46,47^, we observed the expressions of FgTri1-GFP and FgTri4-GFP in PH-1 and GARP complex gene mutants by fluorescence confocal microscopy. The results showed that FgTri4- and FgTri1-labeled toxisomes were almost completely undetectable in the Δ*Fgvps51*, Δ*Fgvps52*, Δ*Fgvps53* and Δ*Fgvps54* mutants but clearly detected in the wild-type PH-1 strain (Fig. 5c, d). These findings indicate that the GARP components FgVps51, FgVps52, FgVps53 and FgVps54 play an indispensable role in DON production and toxin biosynthesis in *F. graminearum*.

### Identification of GARP complex-interacting proteins

To delve into the potential mechanisms by which the GARP complex regulates the pathogenicity of *F. graminearum*, we conducted an affinity capture-mass spectrometry assay to identify GARP complex-interacting proteins. As previously established, FgVps51 and FgVps53 are core components of the GARP complex. Accordingly, vectors expressing FgVps51-GFP and FgVps53-GFP, controlled by their respective natural promoters, were constructed and transformed into wild-type protoplasts to generate FgVps51-GFP- and FgVps53-GFP-expressing strains. Total protein samples from these strains were extracted, and GFP-specific beads were used to pull down the subunits and their interacting proteins, and subsequently subjected to gel electrophoresis (SDS-PAGE). The isolated proteins were further identified using liquid chromatography-mass spectrometry (LC-MS) (Fig. S10a). The results indicated that a total of 1864 and 2506 proteins were pulled from FgVps51-GFP and FgVps53-GFP baits, while the controls PH-1-GFP and PH-1 only pulled 140 and 12 proteins, respectively (Fig. S10b). Further analysis revealed that 1728 proteins were to FgVps51-GFP and FgVps53-GFP (Fig. S10c). To refine this list and identify high-confidence precipitates, peptide candidates with 20% coverage and above were selected. Following this rule, as shown in (Fig. S10d, e), 598 proteins were selected for FgVps51-GFP while 914 proteins were selected for FgVps53-GFP. Of these proteins, 461 were commonly captured from FgVps51-GFP and FgVps53-GFP pulldown, excluding those enriched by PH-1-GFP and PH-1 (Fig. S10f).

To elucidate the biological functions of the putative interacting proteins of FgVps51-GFP and FgVps53-GFP, we conducted Gene Ontology (GO) enrichment analysis on the 461 proteins and identified 20 biological processes (BP) and 43 cellular components (CC). The top fifteen results in each category were sorted after evaluating the Log of their p-values, as a p-value determines the result of a correlation test^48^. For BP, the top-ranking processes include translation, retrograde transport, endosome to Golgi, vesicle-mediated transport, acetyl-CoA biosynthetic process from pyruvate, intracellular protein transport, formation of cytoplasmic translation initiation complex, Arp2/3 complex-mediated actin nucleation, actin cortical patch organization, regulation of actin filament polymerization, tricarboxylic acid cycle, methionine biosynthetic process, ER to Golgi vesicle-mediated transport, cell redox homeostasis, and COPII-coated vesicle budding. For CC, the top-ranking categories include cytosol, ribosome, GARP complex, eukaryotic 48S preinitiation complex, Golgi membrane, retromer, cargo-selective complex, retromer complex, nuclear envelope, endosome, vacuolar proton-transporting V-type ATPase, V1 domain, multi-eIF complex, COPII vesicle coat, cytoplasmic vesicle, and mitochondrial pyruvate dehydrogenase complex, septin complex. Interestingly, the top-ranking biological processes revealed four processes (2, 3, 11, and 15) that are related to vesicle transport (Fig. S11a). Further analysis showed that these proteins were mainly enriched in categories 3, 5, 6, 7, 8, 9, and 11 (Fig. S11b).

### GARP complex interaction with retromer complex

Intriguingly, the Biological Process (BP) enrichment analysis revealed that the second-highest ranked processes included retrograde transport, endosome to Golgi. Subsequent cell enrichment analysis demonstrated that proteins associated with this pathway were predominantly enriched in GARP complexes, retromer complexes, and retromer cargo-selective complexes (Fig. S11). Given that the GARP complex primarily regulates the tethering process of vesicles on the TGN during retrograde transport from endosomes to the TGN^15,20^ and the retromer complex facilitates the transport of various transmembrane proteins from endosomes to the TGN^49,50^, we posit that GARP may play a crucial role in regulating the retromer-mediated transport from endosomes to the TGN. To investigate this possibility, the GARP components FgVps51-mCherry and FgVps53-mCherry were co-expressed with a core subunit of the retromer complex, FgVps35-GFP. Their intracellular localizations were examined by laser scanning confocal microscopy, revealing clear co-localization between FgVps35-GFP and FgVps51-mCherry (Fig. 6a), as well as between FgVps35-GFP and FgVps53-mCherry (Fig. 6b). To further validate their dynamic interaction, Bimolecular Fluorescence Complementation (BiFC) assay was conducted, which confirmed interactions between the retromer subunit FgVps35 and the GARP components FgVps51 and FgVps53 (Fig. 6c, d). The retromer complex, a pentamer crucial for fungal pathogenicity and development, consists of two subcomplexes: a trimer of cargo selective complex (FgVps26-FgVps29-FgVps35) and a dimer of sorting nexins (FgVps5-FgVps17)^33,51,52^. However, the interaction between the components of the GARP complex and the retromer complex remains unclear. In our analysis of direct interaction, five retromer components were respectively cloned into the vector pGADT7 (AD), and the four GARP components cloned into the vector pGBKT7 (BD). Various combinations of these retromer and GARP subunit constructs were co-transformed into *S. cerevisiae* AH109 for Yeast Two-Hybrid (Y2H) assays. Our findings revealed that the GARP complex subunits, FgVps51, FgVps52 and FgVps53, directly interact with the retromer complex subunits FgVps35, FgVps17 and FgVps5 (Fig. 6e, f). These results suggest a direct interaction between the GARP complex and the retromer complex in *F. graminarium*.

**Fig. 6.**
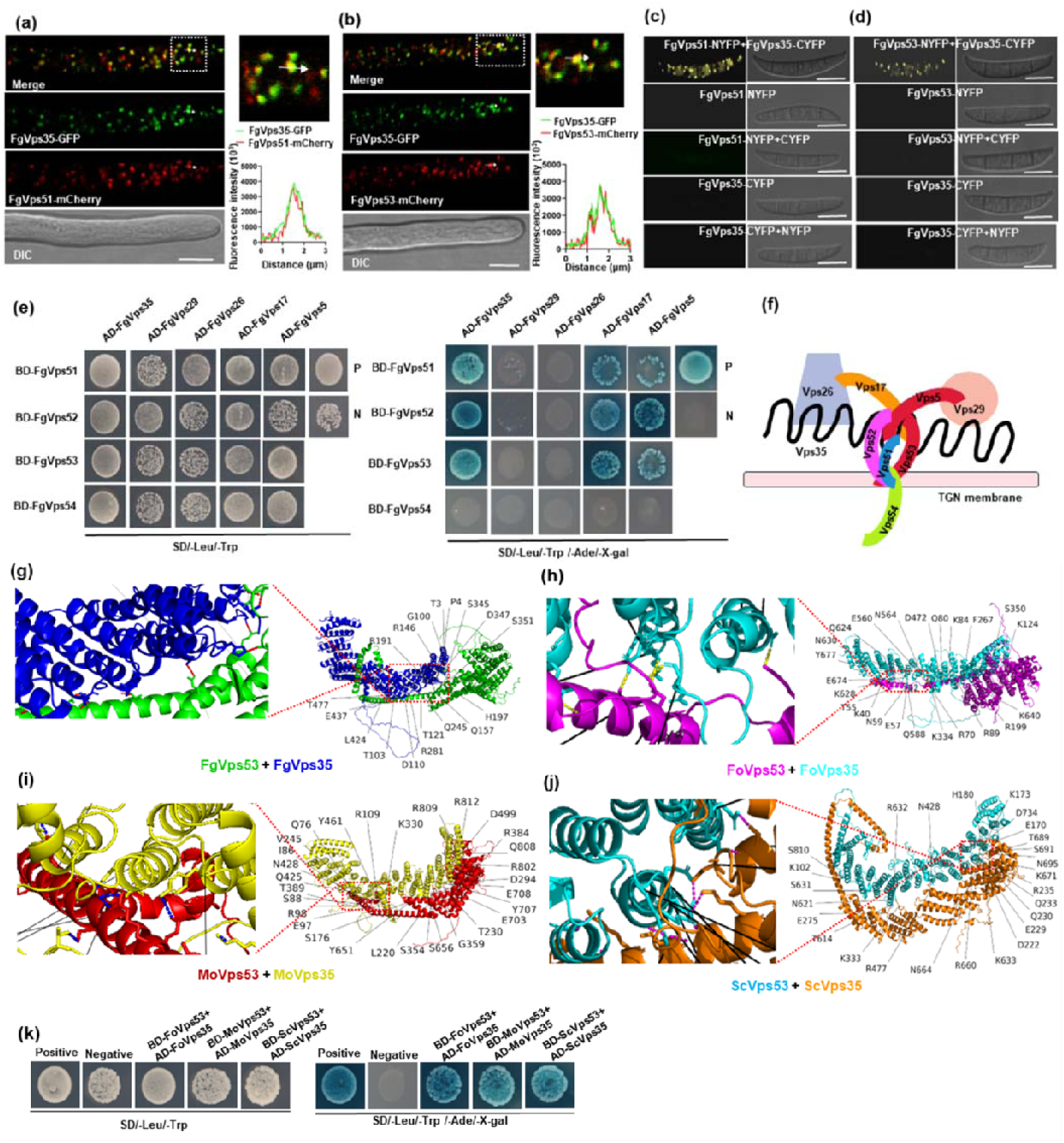
Interaction of GARP with components of the retromer complex. (a) FgVps51-mCherry colocalizes with FgVps35-GFP in *F. graminearum* hyphae. (b) FgVps53-mCherry colocalizes with FgVps35-GFP in the hyphae. White arrowheads indicate the colocalization. Bar=10 µm. (c) Bimolecular fluorescence complementation (BiFC) assay confirming positive interaction between FgVps51-NYFP and FgVps35-CYFP *in vivo*. (d) BiFC assay confirming the interaction of FgVps53 with FgVps35. Bar=10 µm. (e) Y2H assay confirming the interactions between the different components of the GARP complex and those of the retromer complex. The interactions of pGBKT7-53/pGADT7-T and pGBKT7-Lam/pGADT7-T were used as positive and negative controls, respectively. Yeast transformants carrying the indicated constructs are screened for growth on selective plates supplemented with SD-Leu-Trp and SD-Leu-Trp-His-Ade, and for α-galactosidase (LacZ) activities. (f) Interaction model for the GARP complex with the retromer complex. (g) Molecular docking of FgVps53 (green) and FgVps35 (blue) proteins in *F. graminarium*. (h) Molecular docking of FoVps53 (magenta) and FoVps35 (cyan) proteins in *Fusarium odoratissimum*. (i) Molecular docking of MoVps53 (red) and MoVps35 (yellow) proteins in *Magnaporthe oryzae*. (j) Molecular docking of ScVps53 (orange) and ScVps35 (cyan) proteins in *Saccharomyces cerevisiae*. Protein structures were predicted with A_LPHA_F_OLD2_. Molecular docking was performed using the FFT (Fast Fourier Transformed) program GRAMM. (k) Y2H assay confirming the interactions of FoVps53 with FoVps35, MoVps53 with MoVps35 and ScVps53 with ScVps35.

To our knowledge, the direct interaction between the retromer complex and the GARP complex has not been functionally characterized in different species. Therefore, we speculate that the interaction between the GARP complex and the retromer complex may be evolutionarily conserved. To explore this hypothesis, we investigated the possibility of interaction between the homologues of FgVps53 and FgVps35 in *Fusarium odoratissimum* (FoVps53 and FoVps35), *Magnaporthe oryzae* (MoVps53 and MoVps35) and *Saccharomyces cerevisiae* (ScVps53 and ScVps35) by molecular docking analysis. The in-silico analysis revealed specific amino acid residues within these proteins engaged in interactions through hydrogen bonds (Fig. 6g, i, h, j). To confirm their direct interaction, Y2H assays were conducted, which support that the retromer subunit Vps35 interacts with GARP components Vps53 in different species such as *F. odoratissimum*, *M. oryzae* and *S. cerevisiae* (Fig. 6k). Collectively, these results indicate that the GARP complex directly interact with the retromer complex and this interaction is conserved, at least across fungal species.

### The GARP complex recruits the retromer complex to the TGN

Based on the results presented above, we hypothesized that the GARP complex could be involved in facilitating retromer-mediated trafficking. To test this hypothesis, we explored the spatiotemporal dynamics of FgVps51 and FgVps53 along with FgVPS35 using time-lapse microscopy. The results revealed that both FgVps51-GFP, FgVps53-GFP, and FgVps35-mCherry exhibited punctate vesicles, moving together within hyphal cells (Video S1,2). Notably, Video S3 demonstrates that FgVps35-GFP primarily behaves as short-distance, fast-moving vesicles. Furthermore, it consistently moves toward the TGN, as evidenced by its continuous trajectory and co-localization with FgSec7-mCherry (a TGN marker protein) depicted in the video. Furthermore, we co-transformed FgSec7-mCherry and FgVps35-GFP into protoplasts of Δ*Fgvps51* and Δ*Fgvps53* mutants and checked for any alterations in the normal localization of FgVps35-GFP. In the Δ*Fgvps51* mutant, the punctate fluorescence of FgVps35-GFP largely lost its co-localization with FgSec7-mCherry fluorescence as opposed to their clear co-localization observed in the wild-type PH-1 strain (Fig. 7a, b). This is similar to what was observed in the Δ*Fgvps53* mutant, in which the co-localization of FgVps35-GFP and FgSec7-mCherry fluorescence is nearly abolished (Fig 7c). Of 10 independent micrographs, 12.47% ± 2.44% of the fluorescence clear co-localization in PH-1. In contrast, only 3.51% ± 1.38% and 3.42% ± 1.92% of the fluorescence was observed to co-localization in Δ*Fgvps51* and Δ*Fgvps53* mutants, respectively (Fig 7d). On the other hand, we similarly co-expressed FgVps51-GFP and FgVps53-GFP with FgSec7-mCherry in the Δ*Fgvps35* mutant, respectively. The results revealed that the localization of FgVps51-GFP was found to be consistent with that of PH-1, showing no significant changes (Fig 1a and 7e, f). Similarly, the localization pattern of FgVps53-GFP exhibited a comparable situation (Fig 4c and 7g, h). These results strongly suggest that the GARP complex is required for the recruitment of the retromer complex to the TGN but not vice versa.

**Fig. 7.**
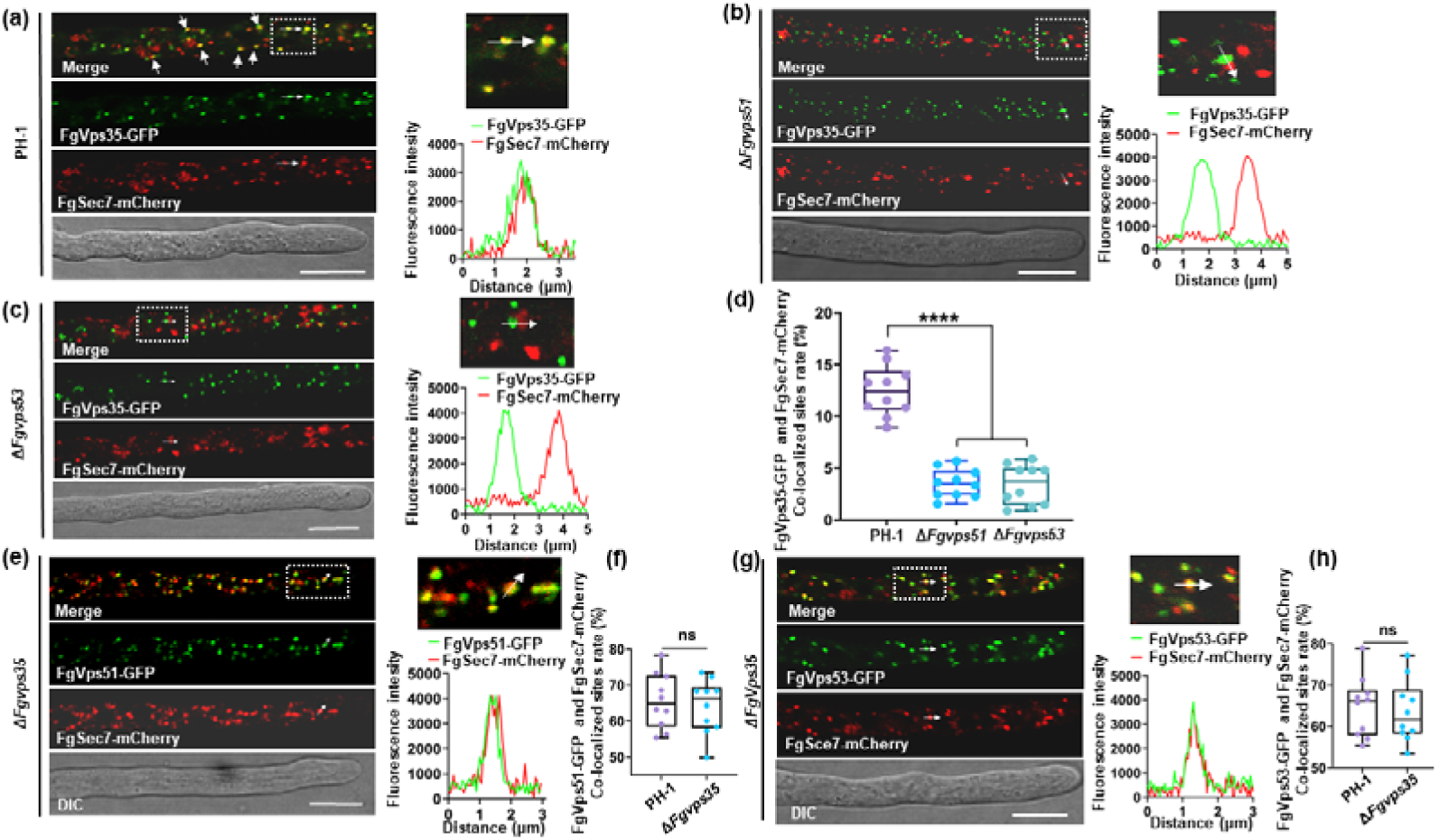
Deletion of *FgVPS51* or *FgVPS53* compromises FgVps35 localization on TGN. (a) In the wild type (PH-1) hyphae, FgVps35-GFP colocalizes with *trans*-Golgi network (TGN, FgSec7-mCherry). White arrows indicate the co-localization sites. (b) In the hyphae of Δ*Fgvps51* mutant, the colocalization of FgVps35-GFP with the TGN marker FgSec7-mCherry was largely lost. White arrows show that the FgVps35-GFP does not co-localize with FgSec7-mCherry at the scanned site. (c) In the Δ*Fgvps53* mutant, there was a partial colocalization of FgVps35-GFP with the TGN. White arrows indicate the incomplete colocalization of FgVps35-GFP with FgSec7-mCherry at the scanned site. (d) Statistical analysis of the co-localization rate of FgVps35-GFP with the TGN in the indicated strains. (e) In Δ*Fgvps35* mutant, FgVps51-GFP co-localize with the TGN. White arrows indicate the co-localization sites. (f) Statistical analysis of the co-localization rates of FgVps51-GFP with the TGN, in the indicated strains. (g) In Δ*Fgvps35* mutant, FgVps53-GFP co-localize with the TGN. White arrows indicate the co-localization sites. (h) Statistical analysis of the co-localization rates of FgVps53-GFP with the TGN, in the indicated strains. Box-plot values represent means of 10 independent experiments. Statistical analysis was processed by two-way ANOVA for multiple comparisons using GraphPad Prism 9 (****p < 0.0001; *ns,* not significant at *P*>0.05).

### FgKex2 and FgSnc1 are cargoes for GARP/retromer-mediated trafficking pathway

Considering the fact that the retromer complex mediates the retrograde trafficking of certain cargo proteins from the plasma membrane/endosomes to the TGN^53,54^, and that the retromer complex requires the GARP for its recruitment to the TGN (Fig. 7), we decided to investigate the potential cargoes involved in the GARP/retromer-mediated vesicle trafficking pathway in *F. graminearum*. We revisited our LC-MS/MS data in which a total of 1856 and 1125 proteins were captured from FgVps51-GFP and FgVps35-GFP pulldown, respectively. To refine this number and identify the high-confidence precipitates, protein candidates that were captured in both sides but absent in the controls, and have a coverage of 20% or more were selected. Using this strategy, a total of 179 proteins were obtained (Fig. S12a, b). To further identify the potential cargo proteins in the GARP/retromer-mediated trafficking pathway, we performed GO enrichment analysis on these 179 proteins, resulting in 13 domain enrichment analysis groups. Interestingly, the top ten domain enrichment groups indicated that the fifth group of these proteins contains transmembrane domains (Fig. S12c). Additionally, the 3D structures of these proteins were predicted by AlphaFold-2. A gene FGSG_09156, encoding a polypeptide of 852 amino acids with transmembrane domains at positions 726-748 (Fig. S13a), and FGSG_08537, encoding a polypeptide of 118 amino acids with putative transmembrane domains at positions 94-116 (Fig. S13b), were identified.

FGSG_09156 is a homologue of *S. cerevisiae* Kex2 protein, a Ca^2+^-dependent serine protease^55^.Therefore, we postulated that FGSG_09156 (FgKex2) could potentially interact with GARP and retromer complexes. To assess this possibility, we conducted a yeast two-hybrid (Y2H) assay and found that FgVps51 and FgVps35 directly interact with FgKex2 (Fig. 8a). Also, BiFC assay further confirmed the interactions between FgVps51/FgVps35 and FgKex2 (Fig. 8b). Furthermore, we hypothesized that FgKex2 acts as a recycling cargo in the GARP/reverse transcriptase-mediated vesicle trafficking pathway from early endosomes to the TGN. To validate our hypothesis, we generated FgKex2-GFP strain and conducted localization experiments. The findings revealed a distinct punctate distribution of FgKex2-GFP across various developmental stages, as illustrated in (Fig. S14). We also observed the predominant presence of FgKex2-GFP in the TGN, with partial localization in endosomes (Fig. 8c, d). Notably, time-lapse microscopy further captured the continuous recycling of FgKex2-GFP to the TGN, evident by its persistent movement towards and co-localization with the TGN marker FgSec7-mCherry, as illustrated in (Video S4). To further substantiate our findings, we used FM4-64 staining and observed that FgKex2-GFP localization was significantly altered in Δ*Fgvps51* and Δ*Fgvps35* mutants, primarily distributed in vacuoles, and failing to localize as punctate structures (Fig. 8e). These data confirmed that the GARP/retromer-mediated vesicle trafficking pathway plays a pivotal role in sorting and transporting FgKex2. In addition, phylogenetic analysis showed that its ortholog and transmembrane (TM) domains are highly conserved in fungi (Fig. S15). However, the systematic studies on Kex2 in filamentous fungi have been lacking. Next, we generated deletion mutants for *FgKEX2* (Δ*Fgkex2*) and confirmed the deletion by Southern blot analysis (Fig. S16). The mutants exhibited defects in growth, conidiation, sexual development, pathogenicity and DON production similar to GARP and retromer complex mutants (Fig. 9). This data supports that FgKex2 is a cargo for GARP/retromer-mediated trafficking pathway, contributing to the development and pathogenicity of *F. graminearum*.

**Fig. 8.**
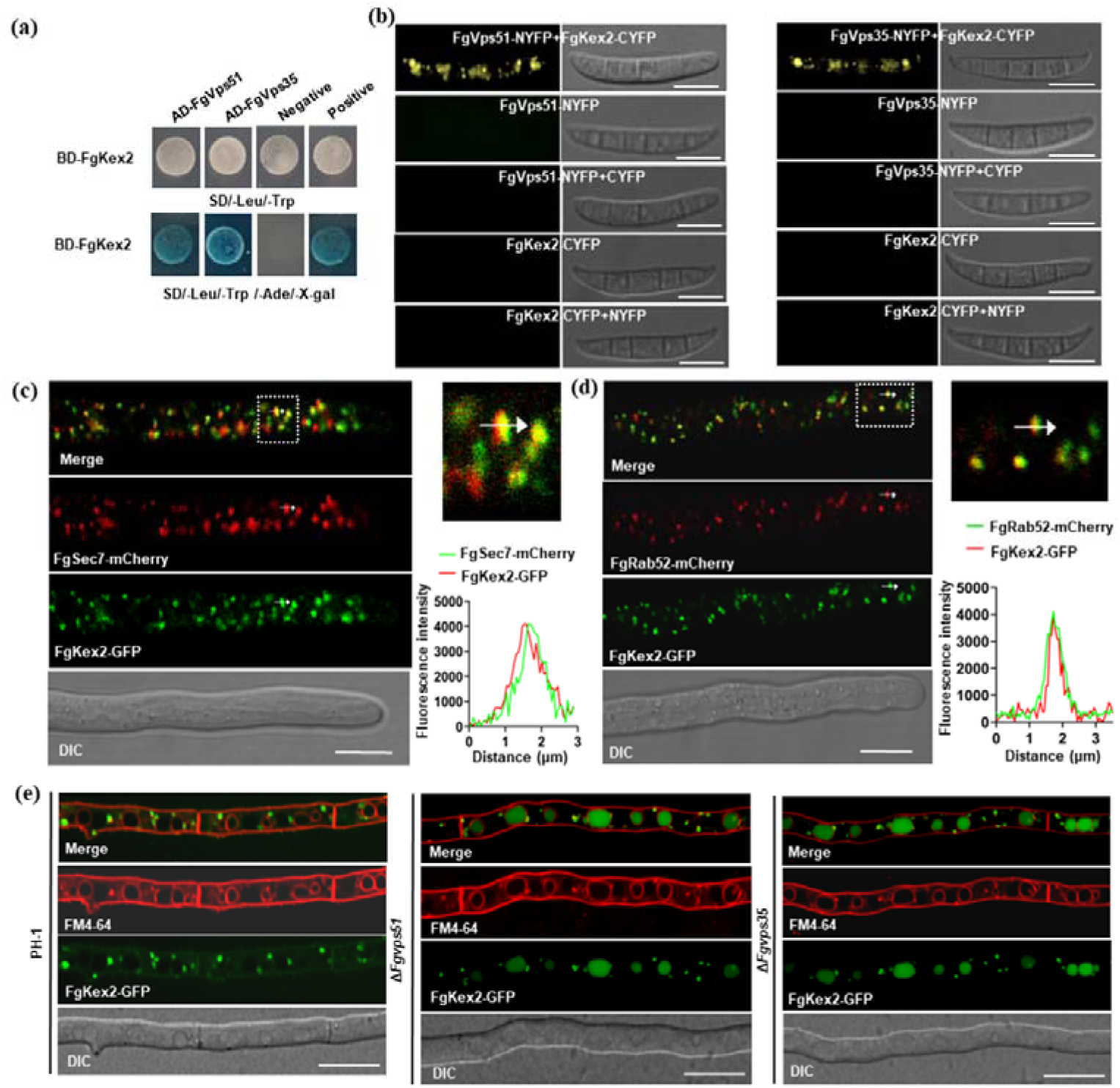
Both FgVps51 and FgVps35 interact with FgKex2 and the association is required for FgKex2 recycling from early endosomes to the Golgi. (a) Y2H assay showing that FgVps51 and FgVps35 interact with FgKex2, respectively, in *F. graminearum*. (b) BiFC assay was used to further confirm the interaction of FgVps51-NYFP and FgVps35-NYFP with FgKex2-CYFP, respectively. The results show that both FgVps35 and FgVps51 interact with FgKex2 *in vivo*. Bar=10 µm (c) FgKex2-GFP partially co-localizes with the *trans*-Golgi network (TGN, FgSec7-mCherry), and the co-localization rate is 61.86 ± 5.52%. White arrows indicate the co-localization sites. Bar=10 µm. (d) FgKex2-GFP partially co-localizes with the early endosome marker mCherry-FgRab52, and the co-localization rate is 37.40 ± 5.52%. White arrows indicate the co-localization sites. Bar=10 µm. (e) Deletion of *FgVPS35* and *FgVPS51* perturbs the transport of FgKex2 from early endosomes to the TGN by mis-sorting FgKex2 to the degradation compartments. Bar=10 µm.

**Fig. 9.**
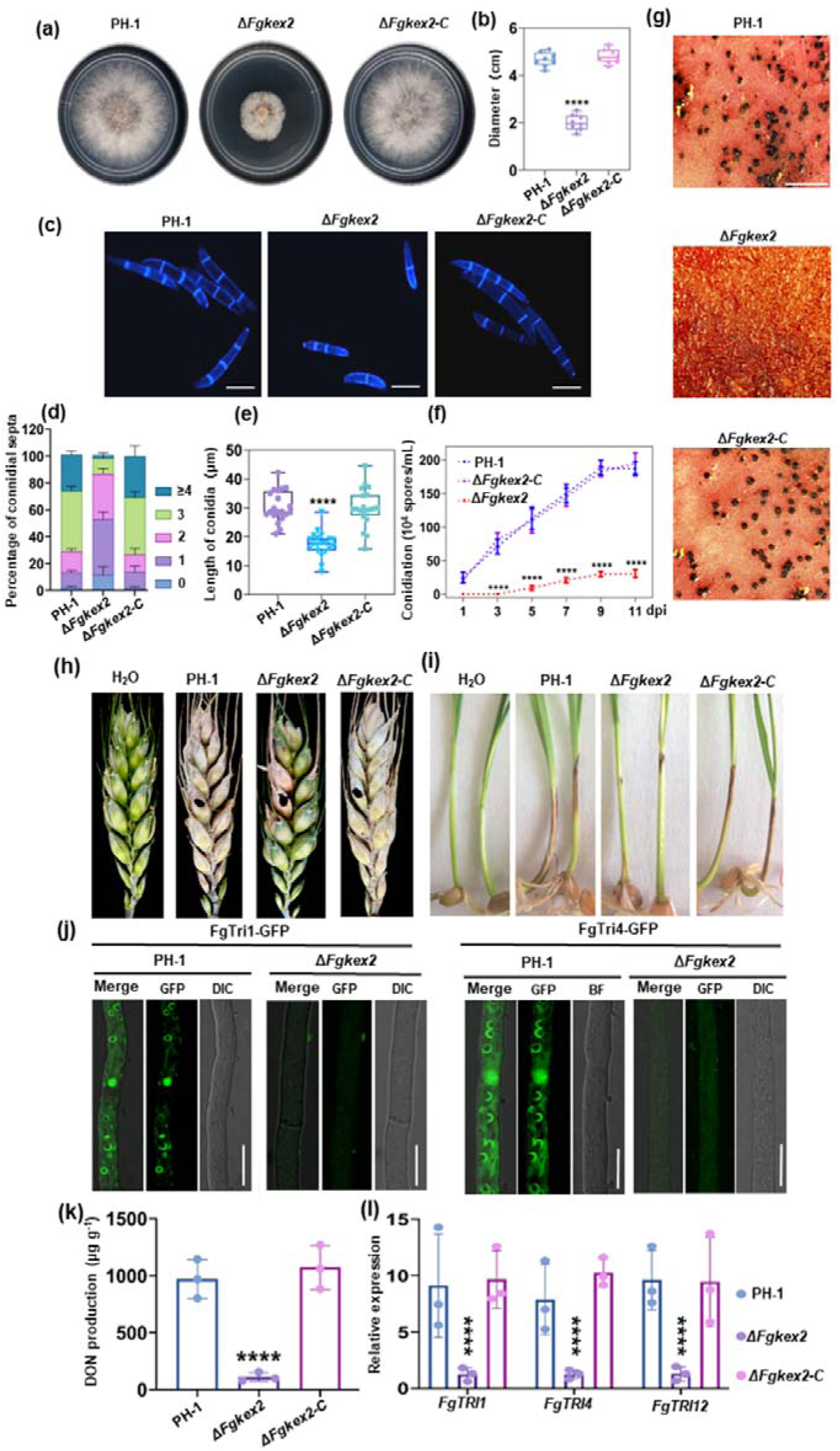
FgKex2 contributes significantly to the vegetative growth, virulence, asexual development and DON production in *F. graminearum.* (a) Vegetative growth of the wild type (PH-1), *FgKEX2* deletion mutant (Δ*Fgkex2*) and complemented strain (Δ*Fgkex2-C*) cultured on complete media (CM) at 28℃ for 3 days. (b) Analyses of the colony diameters of PH-1, Δ*FgKex2* and Δ*FgKex2-C* cultured on CM at 3 dpi. (c) The conidial morphology and number of septa for PH-1, Δ*Fgkex2* and Δ*fgKex2-C* strains stained with 10 µg/mL calcofluor white (CFW) before microscopy. Bar=10 µm. (d) Statistical analysis of the number of septa in the conidia produced by the indicated strains. (e) Analyses of the lengths of the conidia from the tested strains. (f) The number of conidia produced by the PH-1, Δ*Fgkex2* and Δ*Fgkex2-C* strains at different time points. The conidia were harvested from liquid CMC at the indicated time points after inoculation. (g) Perithecial production potential of the indicated strains after growth on carrot agar for two weeks. Bar=1 mm. (h) Pathogenicity of the various strains on flowering wheat heads. The flowering wheat heads were inoculated with mycelial plugs from the indicated strains and the number of diseased spikelets per wheat head was measured at 14 dpi. (i) Pathogenicity of the tested strains to wheat coleoptiles at 7 dpi. (j) Detection of Tri1-GFP and Tri4-GFP fluorescence signals in the tested strains. Bar =10 μm. (k) Analyses of the level of DON produced by the tested strains in 7-day-old TBI cultures. (l) Bar charts showing the relative transcript abundance of TRI genes in the indicated strains detected by quantitative real-time PCR (qPCR). The values shown are means of independent experiments. Statistical analysis was processed by two-way ANOVA for multiple comparisons using GraphPad Prism 9 (∗∗∗∗p < 0.0001).

Furthermore, FGSG_08537, similar to yeast Snc1, was identified. In yeast, Snc1 regulates the fusion of secretory vesicles and the plasma membrane ^56^. In *M. oryzae*, the retromer complex recognizes MoSnc1, promoting its transport to the plasma membrane to regulate vesicle fusion ^32^. However, the functional connection between FgSnc1 and the GARP/retromer-mediated vesicle-trafficking pathway has not been previously established. To address this gap, we developed co-expression strains incorporating GFP-FgSnc1 with FgVps51-mCherry and FgSnc1-GFP with FgVps35-mCherry. Through live cell imaging, we observed significant co-localized punctate structures of GFP-FgSnc1 with FgVps51-mCherry and FgVps35-mCherry in vegetative hyphae and conidia, respectively (Fig. S17). These findings provide conclusive evidence that FgSnc1 indeed interacts with the GARP and retromer components in *F. graminearum*. Our previous research found that in *F.graminearum*, FgSnc1 is mainly located in endosomes, plasma membranes, septa and hyphal apex and is critical for polar hyphal growth and pathogenicity^34^. Moreover, time-lapse microscopy has unveiled the remarkable ability of GFP-FgSnc1 to undergo continuous transport from the TGN toward the hyphal tip, a pivotal mechanism driving polar hyphal growth (Video S5). Therefore, we hypothesize that the GARP/retromer-mediated vesicle trafficking pathway assumes a vital role in facilitating the transport dynamics of GFP-FgSnc1. To validate our hypothesis, FM4-64 staining results demonstrated that the deletion of FgVps51 and FgVps35 impeded the transport of FgSnc1 to plasma membrane structures, septa, and hyphal apex. Instead, FgSnc1 was misdirected to the vacuolar degradation pathway (Fig. S18). These findings strongly suggest the participation of GARP/retromer-mediated vesicle trafficking pathways in the sorting and transport of FgSnc1, which is critical for polar growth and virulence of *F. graminearum*.

## Discussion

Understanding the intricate molecular mechanisms that govern retrograde transport from endosomes to the trans-Golgi network (TGN) is paramount importance. The implications of incorrect sorting and transshipment of cargo cannot be overstated, as it plays a pivotal role in the onset of various illnesses, such as Parkinson’sdisease^57,58^. Hence, it is imperative to employ transport mechanisms, such as the retromer complex, to facilitate the conveyance of membrane proteins from the endosome to both the TGN and the plasma membrane^59^. Notably, our preliminary experimental findings highlight a substantial impact on the development and virulence of *Fusarium graminearum* and *Magnaporthe oryzae* due to the deficiency of the retromer^32,33,60^. Nevertheless, the current understanding of how the retromer complex precisely segregates cargo proteins from endosomes and orchestrates their recycling to the TGN, thereby regulating the development and virulence of fungi, remains elusive. In our research, we have uncovered a novel interaction between the GARP complex and the retromer complex. Additionally, we have substantiated that the GARP complex located on the TGN has the capability to recruit the retromer complex from endosomes, thereby successfully establishing the GARP/retromer transport pathway. Notably, we have pinpointed the pivotal role of the GARP/retromer transport pathway as a crucial coordinator in the sorting and recycling of FgKex2 and FgSnc1 from endosomes to the TGN, effectively shielding them from vacuolar degradation. This process holds profound implications for the growth, development, and pathogenicity of *F. graminearum* (Fig. 10).

**Fig. 10.**
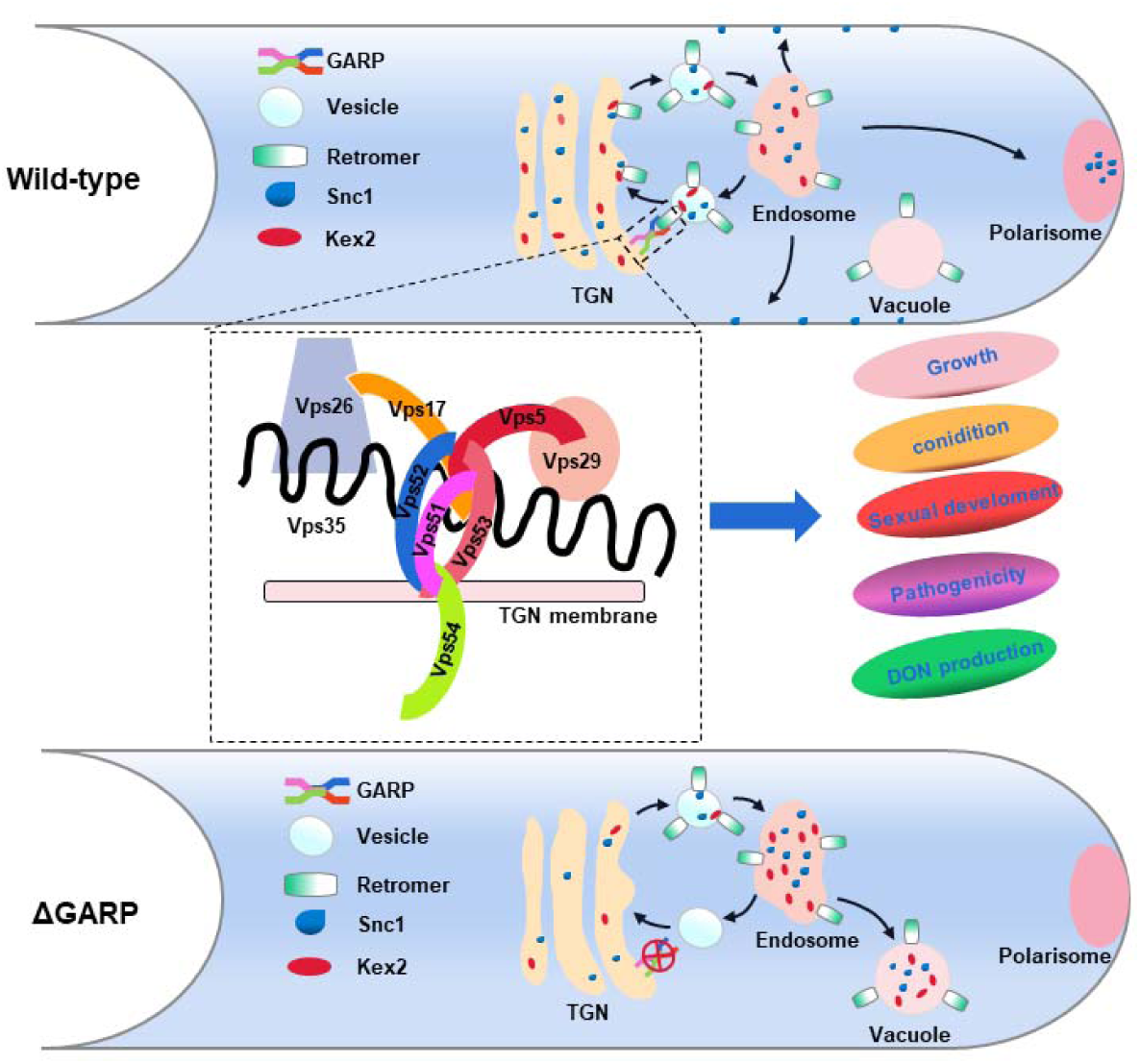
A proposed model for GARP and retromer complex module-mediated vesicle-trafficking pathway and functions in *F. graminearum*. In the wild-type PH-1, the Golgi-associated retrograde protein (GARP) complex regulates retromer-mediated vesicles retrograde transport from early endosomes (EE) to the TGN. Several cargo proteins, like Snc1, on the endosome are recognized by the retromer complex, and then recycled through the GARP-retromer module-mediated vesicular transport pathway to the TGN, which functions in early endocytic recycling, including EE to plasma membrane and EE to hyphal tips. The cargo protein Kex2 is also recycled from early endosomes to the Golgi apparatus via the GARP-retromer module-mediated vesicle trafficking pathway. In the GARP complex null mutant, vesicular retrograde trafficking pathway is greatly impaired, leading to accumulation of Snc1 and Kex2 in the EE. Meanwhile, Snc1 and Kex2 may also be transported to the vacuole and this results in the vacuolar degradation of the proteins. The dashed box represents the interaction of the retromer and GARP complexes. Vps5, Vps52 and Vps53 act as the central components of the GARP complex. They interact with Vps35, Vps17, and Vps5 subunits of retromer complex. The function of the GARP complex in recruiting retromer complex to the TGN is to mediate endosome-to-TGN retrieval, sorting, and transport of cargo proteins. Loss of GARP or retromer-mediated vesicle-trafficking pathway in *F. graminearum* led to impairment of hyphal growth, conidiation, DON production and sexual development as well as pathogenicity.

The conserved presence of the GARP complex across diverse species underscores its deep-rooted evolutionary heritage within the realm of eukaryotes (Figs S1, S4, S5, S6). Recent studies have been accumulating evidence that, while all GARP subunits are non-essential for the survival of yeast^20^, their deletion results in a range of significant outcomes. These consequences encompass heightened protein secretion, disrupted autophagy patterns, and anomalies in vacuolar morphology ^15^. Beyond yeast, the removal of GARP has been linked to infertility in worms^61^, reduced lifespans in flies^62^, and embryonic lethality in mice ^63^. Nevertheless, the role of the GARP complex in pathogenic fungi remains inadequately explored through systematic studies. Hence, deletion mutants of GARP components were generated. The loss of growth, development, and pathogenicity in *F. graminearum* resulting from the deletion of FgVps51, FgVps52, FgVps53, and FgVps54 (Figs 2, 3, 5). Remarkably, deletion of any one of these subunits results in similar phenotypic defects in *F. graminearum*, suggesting the indispensable role of each subunit for the normal functioning of the complex. Additionally, our yeast two-hybrid experiments further demonstrated that these GARP subunits interact to form a tetrameric complex (Fig. 3a), aligning with findings in yeast models^64^. In recent years, mounting evidence has emphasized the critical role of endomembrane trafficking and associated organelles in maintaining physiological processes and pathogenicity in filamentous fungi^32,34^. In yeast, the GARP complex functions as a tethering factor on TGN, ensuring accurate sorting and transportation of cargo proteins^14^. In our study, we made a noteworthy discovery that the subunits of the GARP complex, including FgVps51-GFP, FgVps52-GFP, FgVps53-GFP, and FgVps54-GFP, are ubiquitously expressed throughout all developmental stages and under varying culture conditions (Figs S3, S8). They are primarily situated within the TGN, with partial distribution in endosomes and vacuoles. (Fig. 1, 4). In mammals, for instance, the GARP complex is situated in the TGN, playing a crucial role in coordinating cholesterol transport^65^. In our study, we employed affinity capture mass spectrometry and bioinformatics analysis to identify proteins interacting with the GARP complex. It is noteworthy that the GO enrichment analysis of biological processes indicates that the top-ranking categories, including numbers 2, 3, 11, and 15, are all related to vesicle transport (Fig. S11a). Hence, the GARP complex likely plays a role in coordinating vesicle transport within the TGN to fulfill its biological functions.

Endosomes serve as crucial protein-sorting hubs in eukaryotic cells following endocytosis. One of their essential functions is to manage the movement of cargo proteins through the cell ^66^. The GARP complex plays an important role in this process by mediating the transport of these vesicles to the TGN and facilitating their fusion with the TGN. This ensures the efficient recycling of the cargo proteins^14^. However, very little is known about the various proteins that depend on the GARP complex for their recycling. Herein, we uncovered over 461 potential GARP-interacting proteins (Fig. S9). The retromer complex, which is known to be involved in cargo sorting and transport from endosomes to the TGN, was identified among these proteins. The retromer complex plays a critical role in preventing unnecessary cargo degradation by efficiently sorting cargoes from endosomes and directing them toward the TGN through the retrograde trafficking pathway^67^. In our research, Through the use of high-resolution time-lapse video microscopy, we have observed that FgVps35-GFP primarily behaves as short-distance, fast-moving vesicles. Furthermore, it consistently moves toward the TGN, as evidenced by its continuous trajectory and co-localization with FgSec7-mCherry (a TGN marker protein) depicted in the video (Video S3). This dynamic process plays a pivotal role in the efficient transport of cargo proteins to the TGN, thereby regulating the development and pathogenicity of *F. graminearum*^33^. However, the precise mechanism by which the retromer complex is recruited to the TGN membrane to facilitate the retrograde transport of cargo proteins is largely mysterious. In our study, we investigated the spatiotemporal dynamics of FgVps51–FgSVps35 and FgVps53–FgSVps35 using time-lapse microscopy. The results revealed that both FgVps35-GFP and FgVps51-mCherry, as well as FgVps35-GFP and FgVps53-mCherry, manifest as punctate vesicles and travel together inside hyphal cells (Video S1.2). In addition, we established the interaction model between the GARP complex and the retromer complex through co-localization, BiFC (Bimolecular Fluorescence Complementation), and yeast two-hybrid assays (Fig. 6f). Furthermore, we found that this interaction is highly conserved across different fungal species (Fig. 6k). Our data revealed that absence of GARP functions disrupted the proper localization of the retromer complex to the TGN (Figs 7b,7c); This observation confirmed that the GARP complex indeed has the capability to recruit the retromer complex, thus facilitating the retrograde transport of cargo proteins from endosomes to the TGN. In summary, we have, for the first time, elucidated the GARP/retromer transport pathway from endosomes to the TGN in detail.

In our previous research, we established the involvement of the retromer complex in vesicle transport in *M. oryzae* and *F. graminearum*, highlighting its significance in the growth, development, and virulence of filamentous fungi^32,33,60^. However, the mechanism by which the GARP complex collaborates with the retromer complex to sort and recycle cargo proteins from endosomes to the TGN, thereby regulating fungal polarity growth and pathogenesis, has remained elusive. One of the extensively studied functions of the GARP complex and retromer complex, respectively, is to facilitate the recycling of transmembrane proteins from endosomes to the TGN, exemplified by the vacuolar protein sorting receptor Vps10p in *Saccharomyces cerevisiae*^20,68^. In our current study, we employed high-resolution interactome analysis of GFP trap-precipitated FgVps51-GFP and FgVps35-GFP. Furthermore, domain GO enrichment analysis unveiled the presence of transmembrane domains in FgKex2 and FgSnc1 (Fig S11d). In addition, utilizing time-lapse microscopy, we observed the spatiotemporal dynamics of FgKex2-GFP, noting its punctate appearance and movement toward the Golgi organelle marker, FgSec7-mCherry, within *F. graminearum* cells (Video S4). Through co-localization, BiFC (Bimolecular Fluorescence Complementation), and yeast two-hybrid experiments, we established that both FgKex2 and FgSnc1 interact with the GARP and retromer complexes (Figs 8a, b and S17). Importantly, the deletion of either the GARP or retromer complex led to the misrouting of FgKex2 and FgSnc1 toward the degradation pathway (Figs 8e and S18). This phenomenon is attributed to the GARP and retromer complex mutants’ inability to correctly recycle the target proteins FgSnc1 and FgKex2 from late endosomes to the TGN, leading to erroneous sorting and transport. This aligns with previous reports that misclassification of FgSnc1 in filamentous fungi severely disrupts the polarized growth of hyphae^32,34^. Moreover, Kex2 protease, which enzymatically cleaves and modifies effector factors in *Ustilago maydis*, plays a pivotal role in the pathogenicity of filamentous fungi^69–71^. Intriguingly, while systematic studies on Kex2 in filamentous fungi have been lacking, our research revealed a relatively conserved presence of Kex2 in various fungal species (Fig. S15). Furthermore, we found that the deletion of FgKex2 exhibited defects in growth, conidiation, sexual development, pathogenicity, and DON production of *F. graminearum*, akin to GARP and retromer complex mutations (Fig. 9). In conclusion, our data strongly suggest the participation of GARP/retromer-mediated vesicle trafficking pathways in the sorting and transport of FgSnc1 and FgKex2, which is critical for growth, development, and pathogenicity of *F. graminearum*.

However, for detailed roles functional mechanism of the GARP complex in vesicles trafficking, further studies need to be conducted to dissect of the link between this protein complex and GTPases, SNAREs and other transport proteins. A critical consideration is whether GARP collaborates with other tethering factors to accomplish its functions. Furthermore, it remains an open question whether GARP possesses other functions beyond its tethering role at the TGN. Bridging these knowledge gaps will deepen our understanding of the mechanism and functions of GARP complex.

## Author contributions

Conceptualization, WZ and YL; methodology, YL and WZ; investigation, YL, XC, JC, HZ, YL, SC, NP, XZ and RS; visualization, HZ, YY, GL, ZW and WZ; Writing – Original Draft, YL, WZ; Writing – Review & Editing, YL, YSA, ZW and WZ; resources and funding acquisition, WZ.

## Acknowledgements

We appreciate Prof. Jie Zhou, Drs LiLi Lin, Jiexiong Hu, Wenqin Fang, as well as Shuai Yang and Dingyang Zhang for their helpful suggestions. This research was supported by the National Natural Science Foundation of China (32272481, 32122071), and the Natural Science Foundation of Fujian Province (2021J06015).

---

**Fig. S1.**
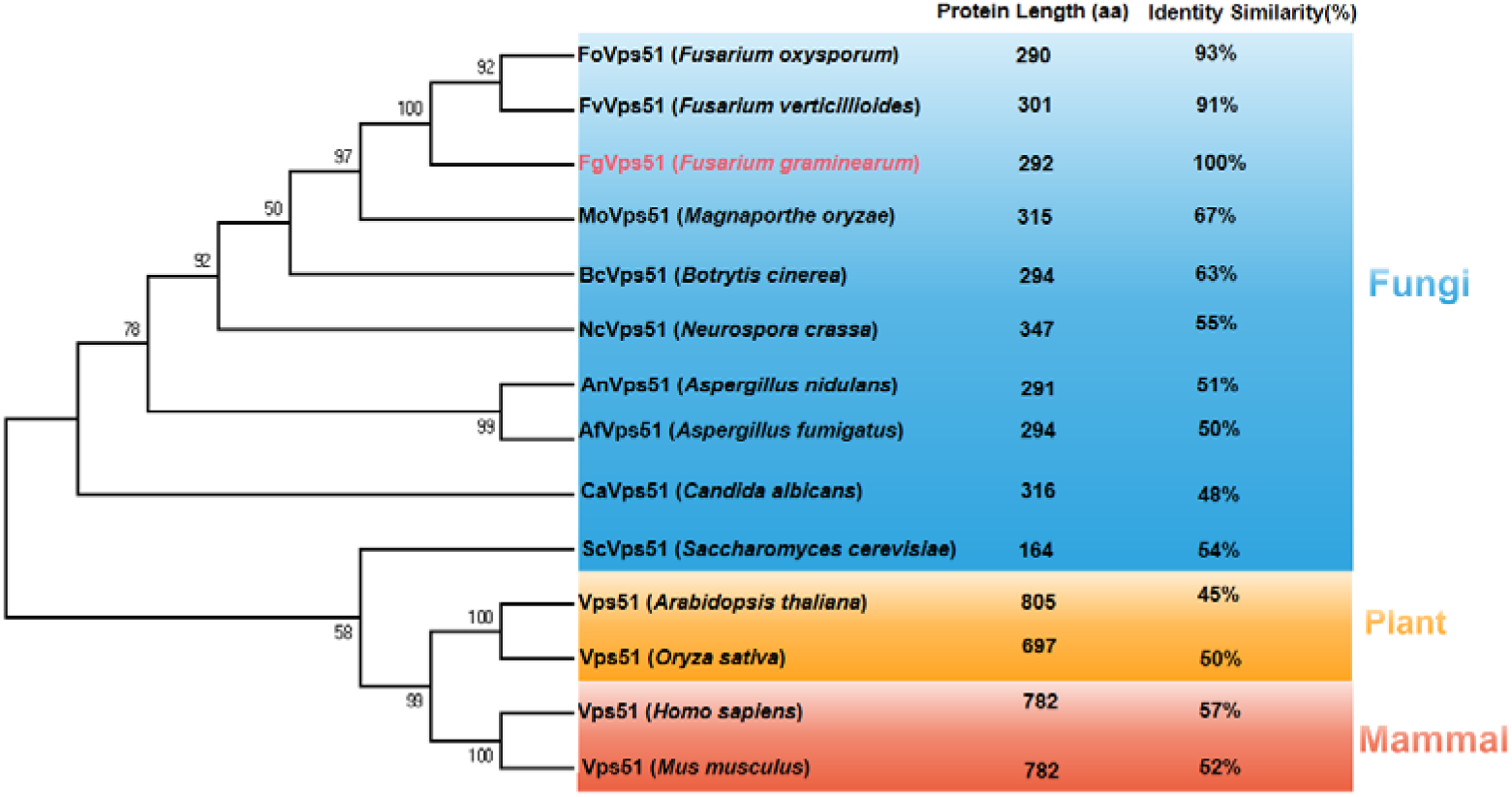
Phylogenetic analysis of putative Vps51 orthologs in fungi, plants and human. The analysis was performed using MEGA7 Program. The GenBank accession numbers (corresponding species names) of the Vps51 orthologs are as follows: KAG7418196.1 (FoVps51 *Fusarium oxysporum*), XP_018749768.1 (FvVps51 *Fusarium verticillioides)*, XP_011324974.1 FgVps53 (*Fusarium graminearum*), XP_003716699.1 (MoVps51 *Magnaporthe oryzae*), XP_024547340.1 (BcVps51 *Botrytis cinerea*), XP_959546.2 (NcVps51 *Neurospora crassa*), XP_660619.1 (AnVps51 *Aspergillus nidulans*), XP_754737.1 (AfVps51 *Aspergillus fumigatus*), XP_719130.2 (CaVps51 *Candida albicans*), NP_012945.1 (ScVps51 *Saccharomyces cerevisiae*), AEE82114.1 (Vps51 *Arabidopsis thaliana*), EEE59892.1 (Vps51 *Oryza sativa*), NP_037397.2 (Vps51 *Homo sapiens*), NP_001074510.1 (Vps51 *Mus musculus*).

**Fig. S2.**
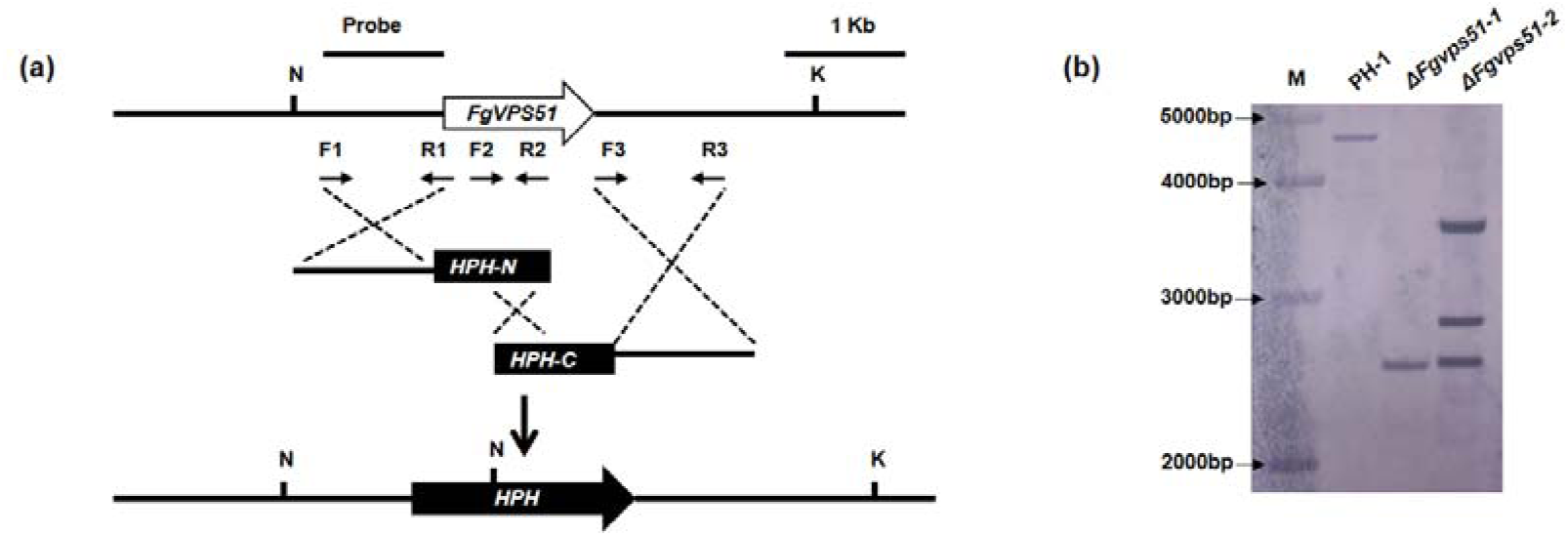
The targeted gene replacement strategy and Southern blot assay. (a) *FgVPS51* targeted gene replacement strategy is shown in the schematic diagram. The primer pairs F1/R1 and F3/R3 were used to generate the gene replacement constructs. Primers F2 and R2 were used for mutant screening and identification. (b) Southern blot confirmation of *FgVPS51* gene deletion. *Nco* l (N)- and *Kpn* l (K)-digested DNAs showed a 4.62 kb band in the PH-1 and a 2.20 kb band in the mutants, confirming that the Δ*Fgvps51-1* generated is the right mutant.

**Fig. S3.**
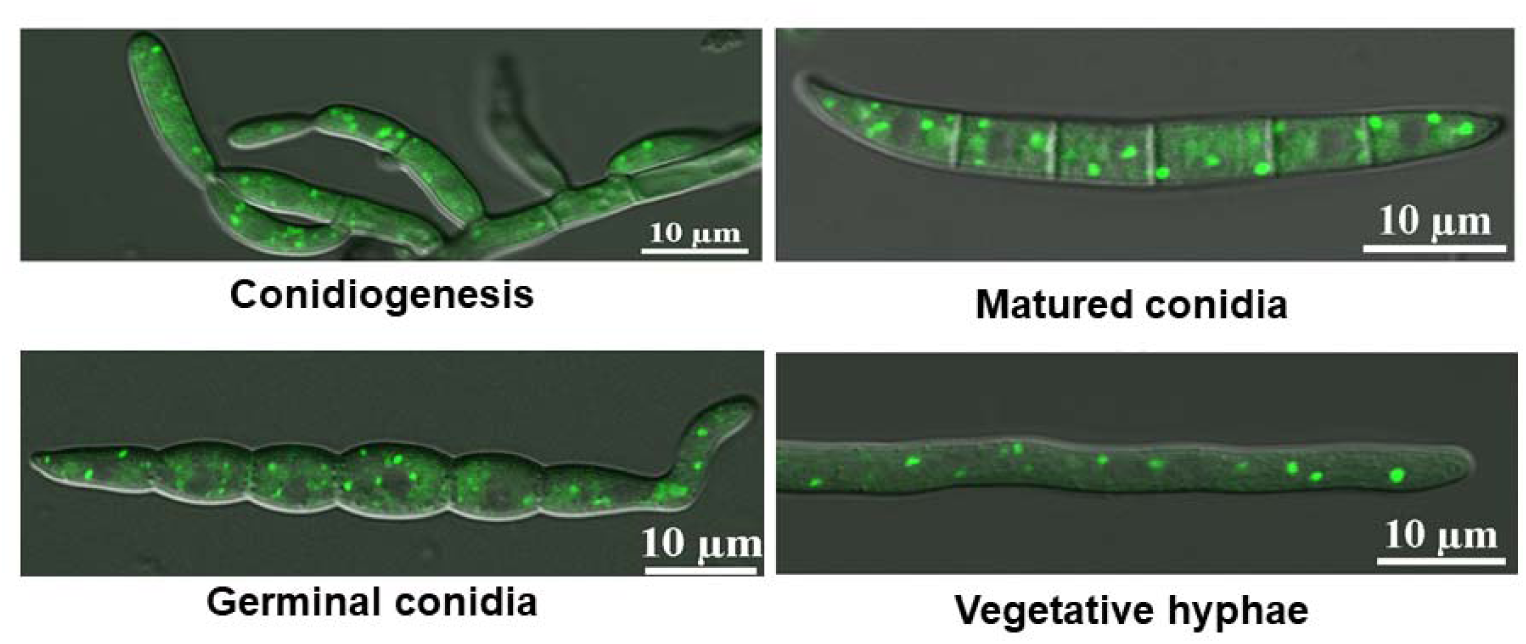
Localization of FgVps51-GFP. The expression of FgVps51-GFP at different developmental stages was captured by Nikon TiE inverted laser scanning confocal microscope system. The fusion protein is localized in the cytoplasm of phialide cells and primary conidia, mature conidia, germinated conidia and vegetative hyphae as small dots.

**Fig. S4.**
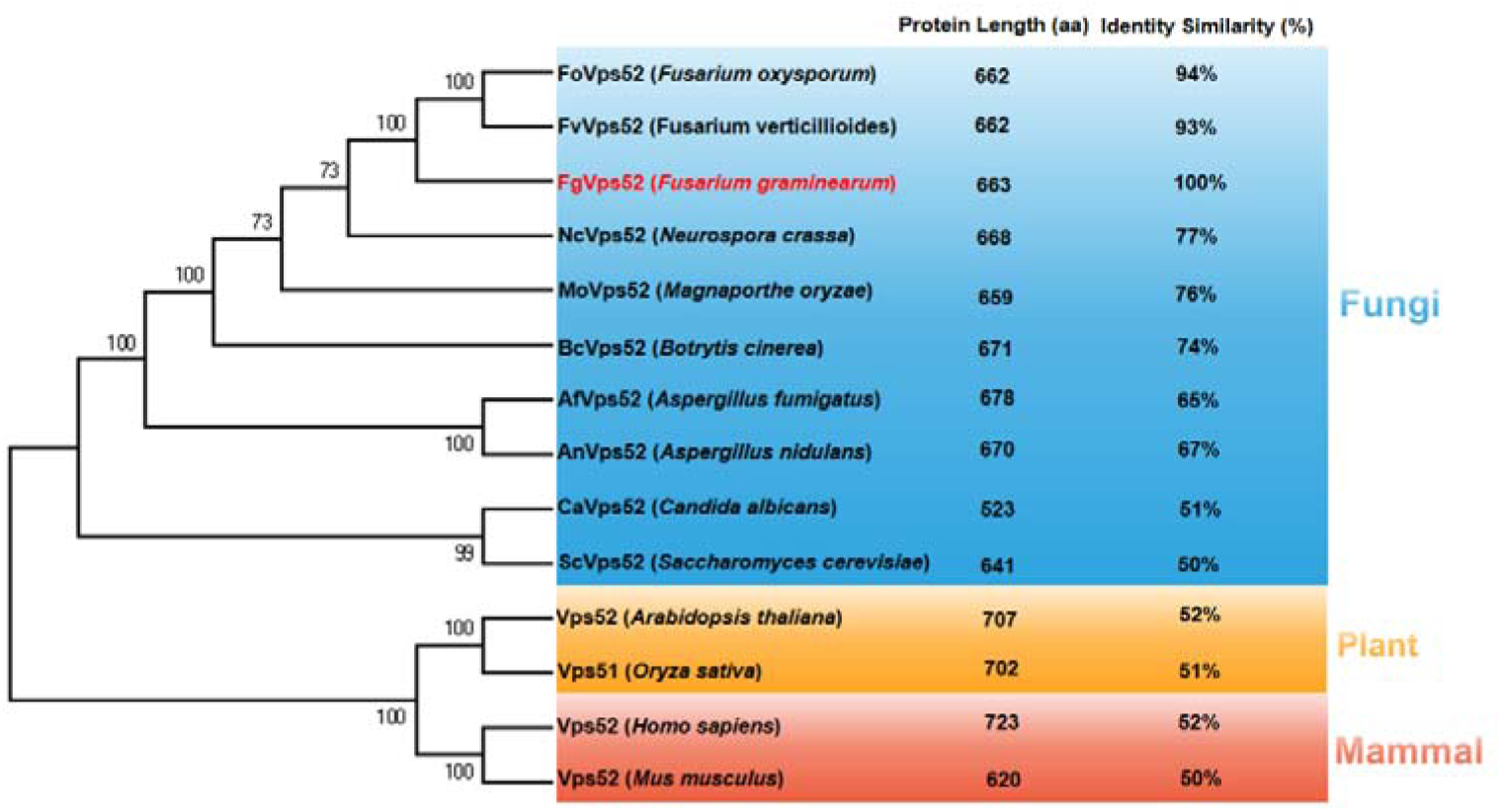
Phylogenetic analysis of putative Vps52 orthologs in fungi, plants and mammal. The analysis was performed using MEGA7 Program. The GenBank accession numbers (corresponding species names) of the Vps51 orthologs are as follows: EXM22288.1 (FoVps52 *Fusarium oxysporum*), XP_018753243.1 (FvVps52 *Fusarium verticillioides*), XP_011317809.1 (FgVps52 *Fusarium graminearum*), XP_961910.2 (NcVps52 *Neurospora crassa*), XP_003711867.1 (MoVps52 *Magnaporthe oryzae*), XP_024549623.1 (BcVps52 *Botrytis cinerea*), XP_750157.2 (AfVps52 *Aspergillus fumigatus*), XP_050467381.1 (AnVps52 *Aspergillus nidulans*), XP_719419.1 (CaVps52 *Candida albicans*), QHB07941.1 (ScVps52 *Saccharomyces cerevisiae*), NP_565015.1 (Vps52 *Arabidopsis thaliana*), XP_015632567.1 (Vps51 *Oryza sativa*), UQL51221.1 (Vps52 *Homo sapiens*), AAH63329.1 (Vps52 *Mus musculus*).

**Fig. S5.**
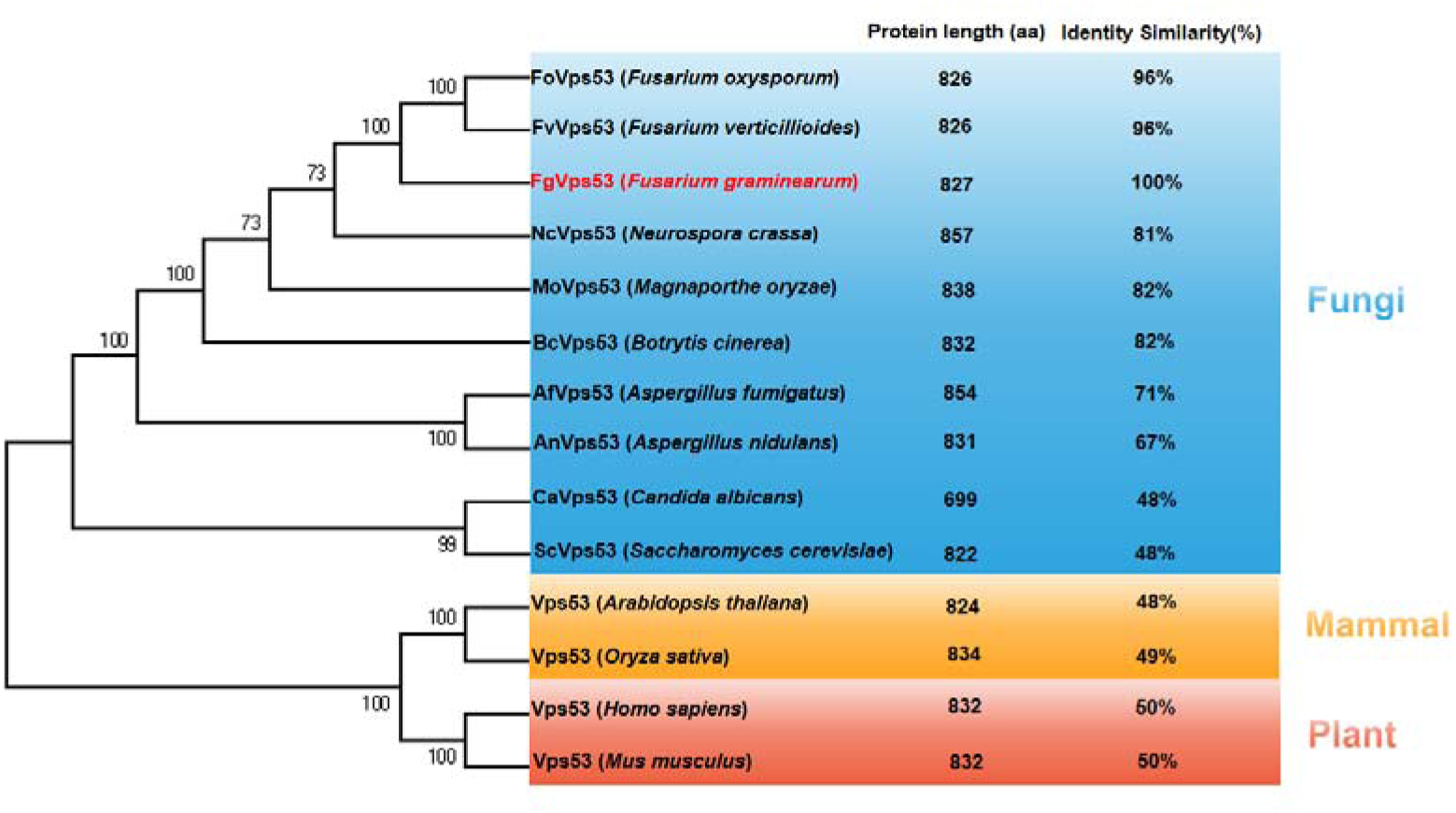
Phylogenetic analysis of putative Vps53 orthologs in fungi, plants and mammal. The analysis was performed using MEGA7 Program. The GenBank accession numbers (corresponding species names) of the Vps53 orthologs are as follows: KAG7420912.1 (FoVps53 *Fusarium oxysporum*), XP_018742921.1 (FvVps53 *Fusarium verticillioides*), XP_011316597.1 (FgVps53 *Fusarium graminearum*), XP_965410.1 (NcVps53 *Neurospora crassa*), XP_003714256.1 (MoVps53 *Magnaporthe oryzae*), XP_024548699.1 (BcVps53 *Botrytis cinerea*), XP_750259.1 (AfVps53 *Aspergillus fumigatus*), XP_660340.1 (AnVps53 *Aspergillus nidulans*), XP_719053.1 (CaVps53 *Candida albicans*), AJV41983.1 (ScVps53 *Saccharomyces cerevisiae*), NP_001322422.1 (Vps53 *Arabidopsis thaliana*), XP_015622127.1 (Vps53 *Oryza sativa*), NP_001121631.1 (Vps53 *Homo sapiens*), NP_080940.2 (Vps53 *Mus musculus*).

**Fig. S6.**
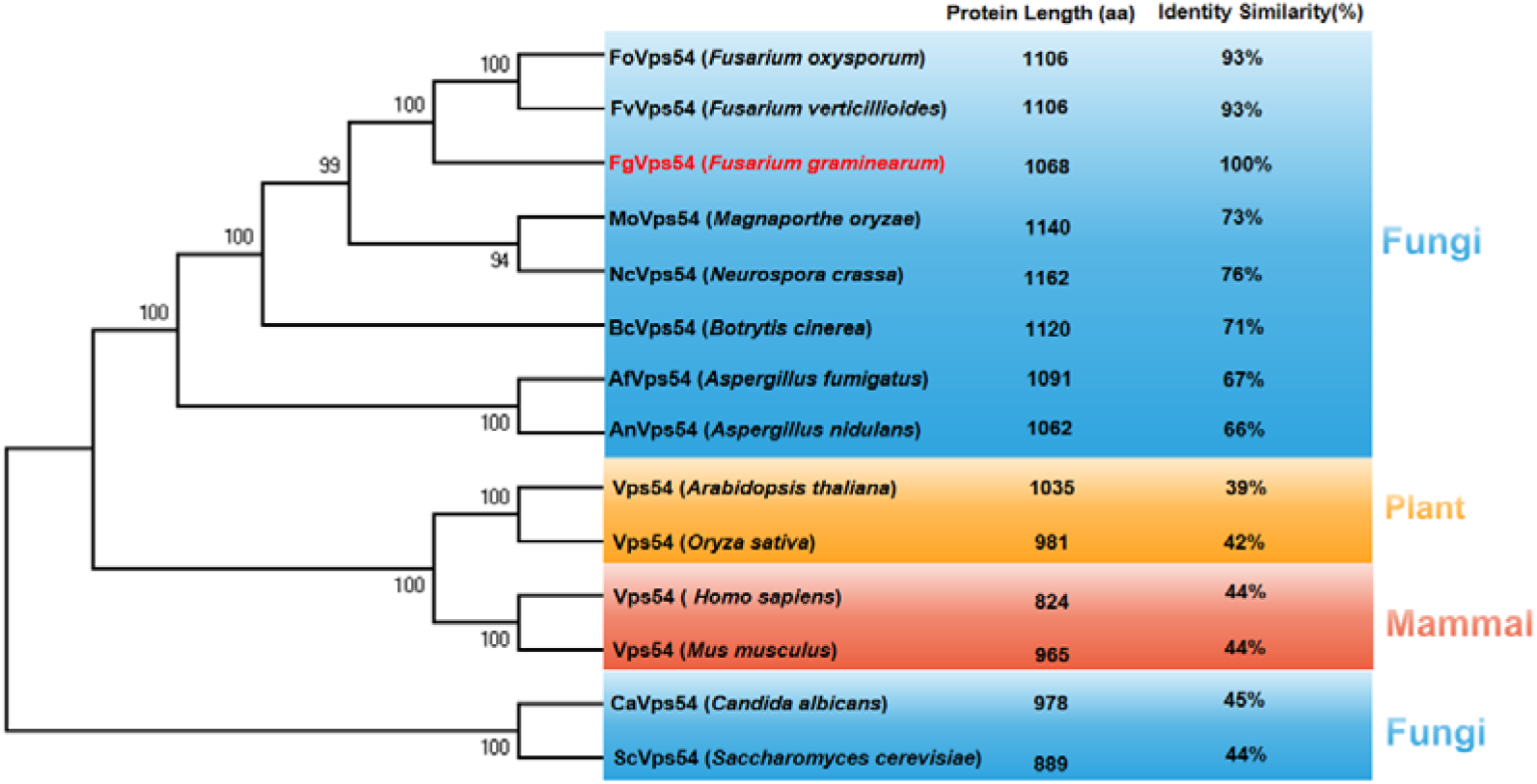
Phylogenetic analysis of putative Vps54 orthologs in fungi, plants and mamml. The analysis was performed using MEGA7 Program. The GenBank accession numbers (corresponding species names) of the Vps54 orthologs are as follows: KAI3577058.1 (FoVps54 *Fusarium oxysporum*), RBQ66322.1 (FvVps54 *Fusarium verticillioides*), XP_011326428.1 (FgVps54 *Fusarium graminearum*), XP_003711076.1 (MoVps54 *Magnaporthe oryzae*), XP_961538.2 (NcVps54 *Neurospora crassa*), XP_024553342.1 (BcVps54 *Botrytis cinerea*), EDP51102.1 (AfVps54 *Aspergillus fumigatus*), XP_681262.1 (AnVps54 *Aspergillus nidulans*), OAO99823.1 (Vps54 *Arabidopsis thaliana*), XP_015636554.1 (Vps54 *Oryza sativa*), AAH41868.1 (Vps54 *Homo sapiens*), NP_001277557.1 (Vps54 *Mus musculus*), KGU15996.1 (CaVps54 *Candida albicans*), CAY78536.1 (ScVps54 *Saccharomyces cerevisiae*).

**Fig. S7.**
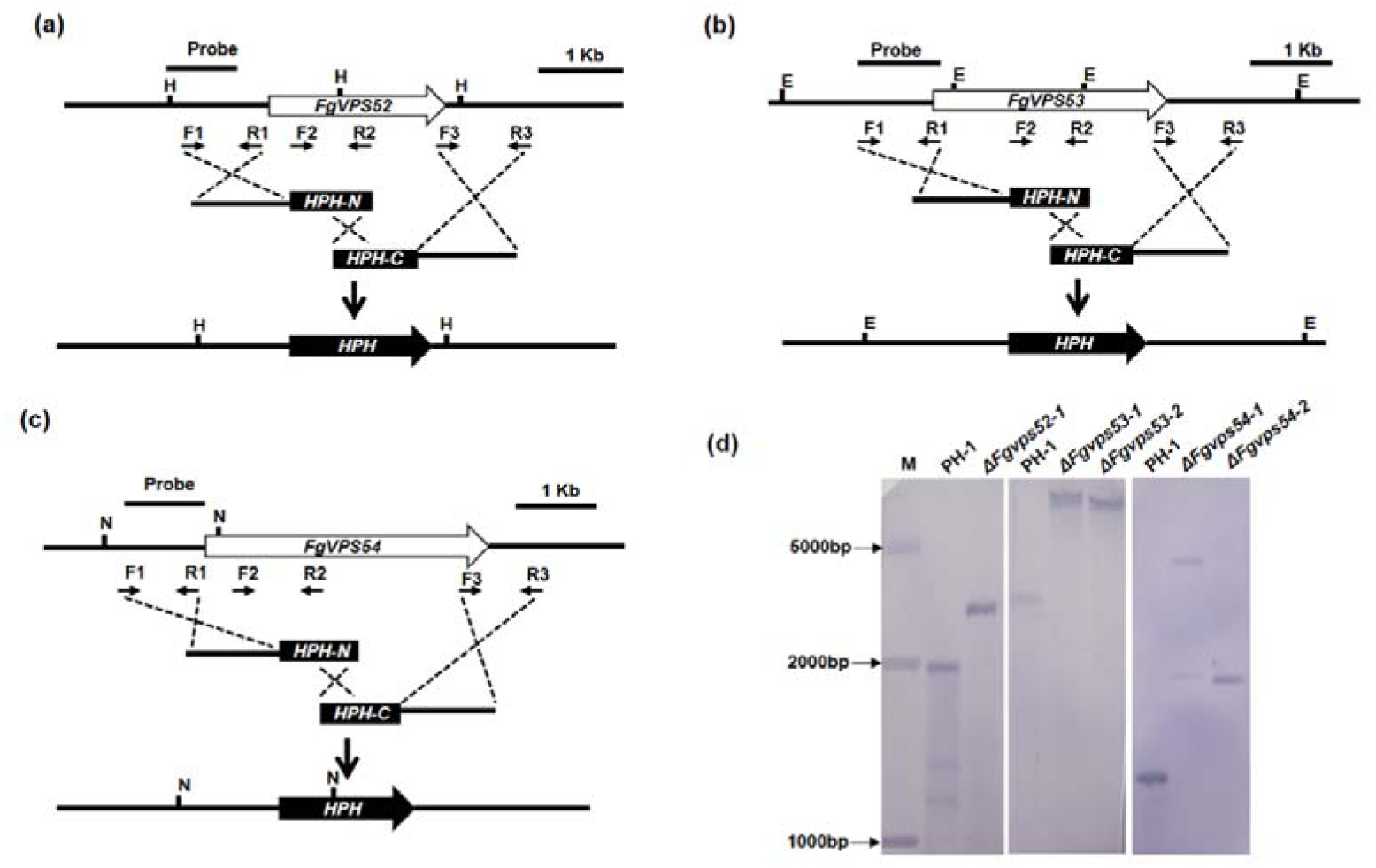
Targeted gene replacement strategy and Southern blot assay. The targeted gene replacement strategies are shown Schematically. The primer pairs F1/R1 and F3/R3 were used to generate the gene replacement constructs. Primers F2 and R2 were used for mutant screening and identification. (a) Targeted gene deletion strategy for *FgVPS52*. *Hin*dlll (H)-digested DNAs showed a 1.94 kb band in PH-1 and a 2.57 kb band in the mutants. (b) Targeted gene deletion strategy for *FgVPS53*. *Eco*Rv (E)-digested DNAs showed a 2.59 kb band in PH-1 and a 6.41 kb band in the mutants. (c) Targeted gene deletion strategy for *FgVPS54*. *Nco*l(N)-digested DNAs showed a 1.35 kb band in PH-1 and a 1.97 kb band in the mutants. (d) Southern blot assay for confirmation of the gene deletions.

**Fig. S8.**
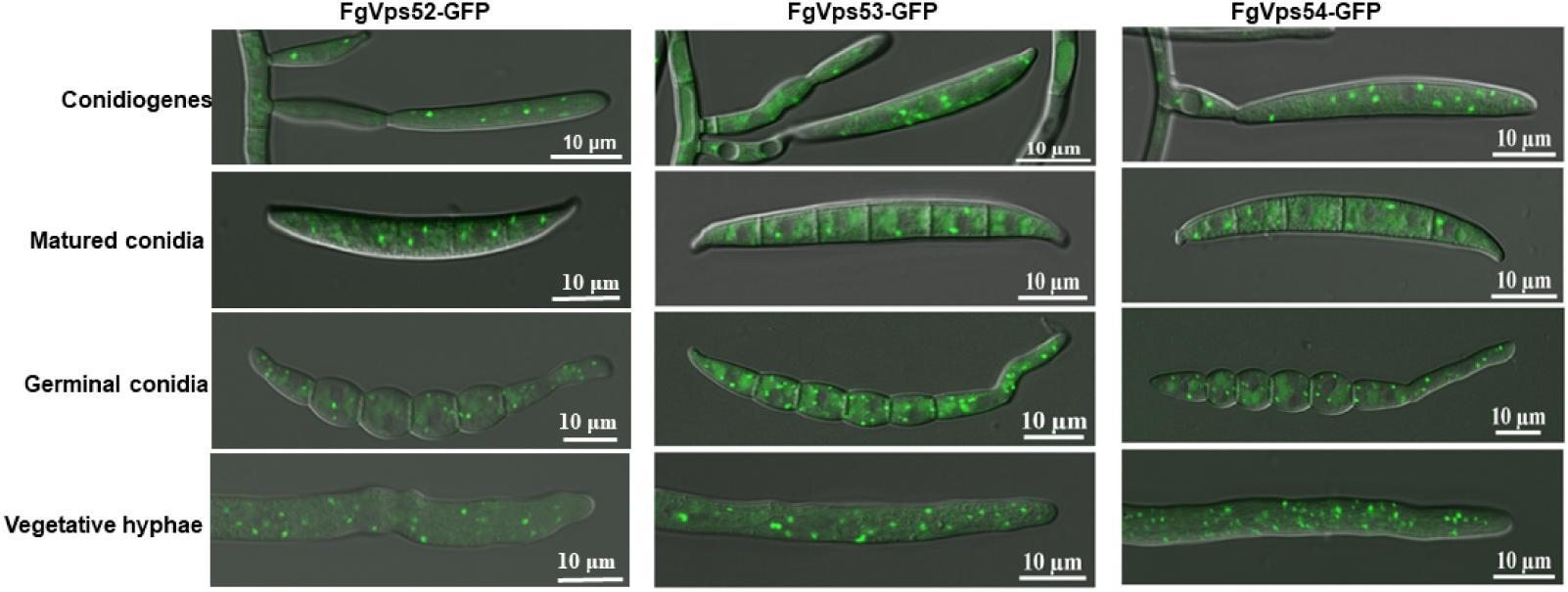
Subcellular localizations of FgVps52-GFP, FgVps53-GFP and FgVps54-GFP. The expression of FgVps52-GFP, FgVps53-GFP and FgVps54-GFP at different developmental stages were captured by Nikon TiE inverted laser scanning confocal microscope system. The fusion proteins are localized in the cytoplasm of phialides cells and primary conidia, mature conidia, germinated conidia and vegetative hyphae as small dots.

**Fig. S9.**
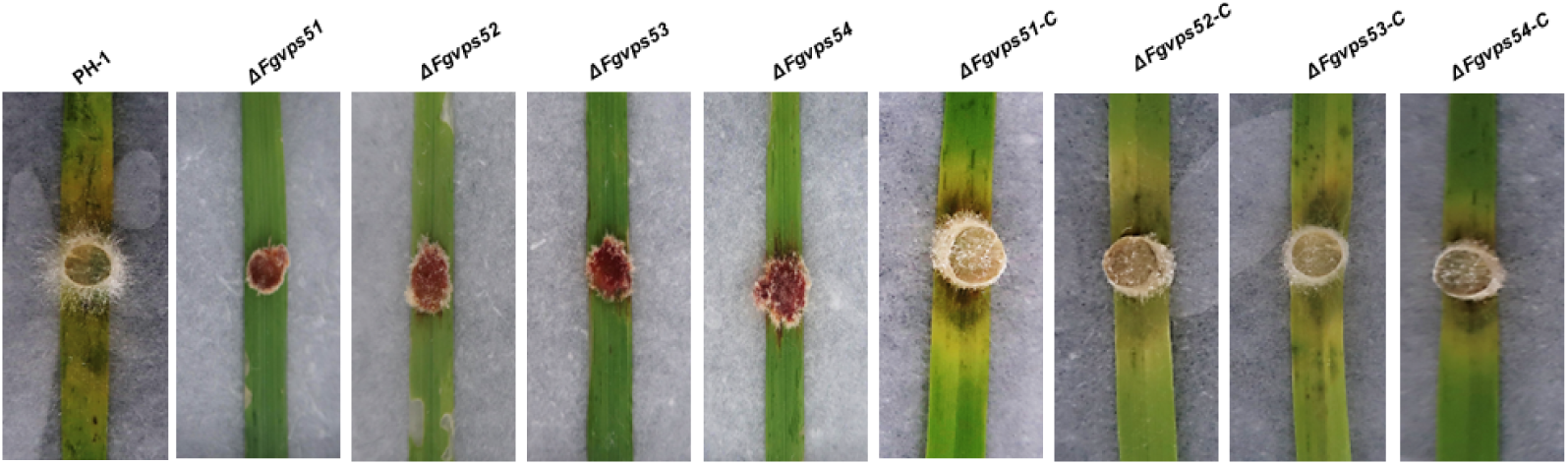
The GARP complex is important for *F. graminearum* virulence to wheat leaves. Mycelial plugs from fresh cultures of the indicated strains were used to infect wheat seedling leaves.

**Fig. S10.**
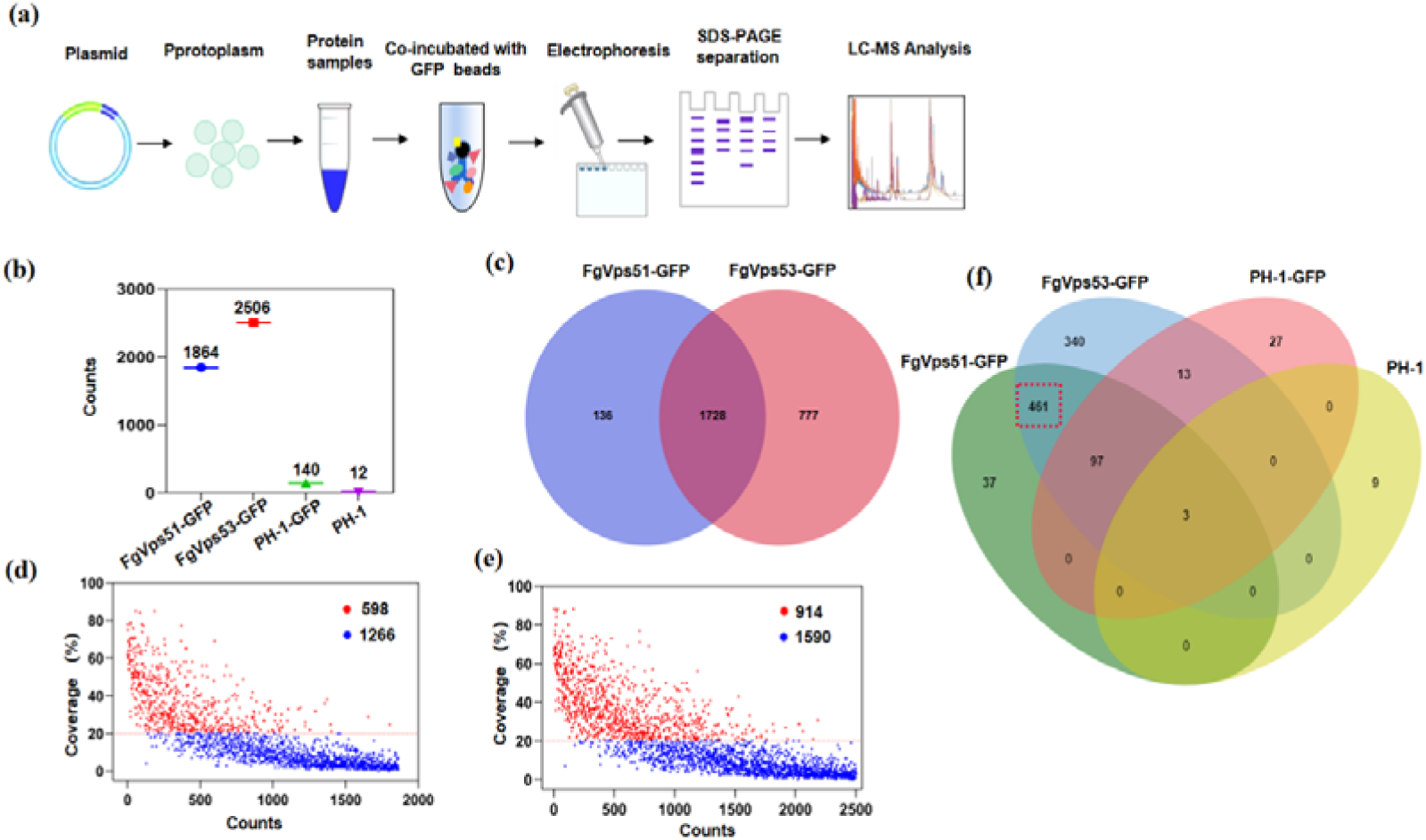
Identification of FgVps51- and FgVps53-interacting proteins. (a) Schematic diagram of FgVps51-GFP and FgVps35-GFP capture-mass spectrometry assay. (b) The number of proteins identified by affinity purification and mass spectrometry. (c) Venn diagram showing the FgVps51- and FgVps53-interacting proteins; 1728 proteins were common to the two biological repeats. (d) 598 FgVps51-interacting partners were identified with 20% coverage and above. (e) 914 FgVps53-interacting partners were identified with 20% coverage and above. (f) In the Venn diagram of the FgVps51-GFP and FgVps53-GFP interaction groups, 461 proteins were specifically identified with high confidence, but not in the PH-1 and PH-1-GFP control groups.

**Fig. S11.**
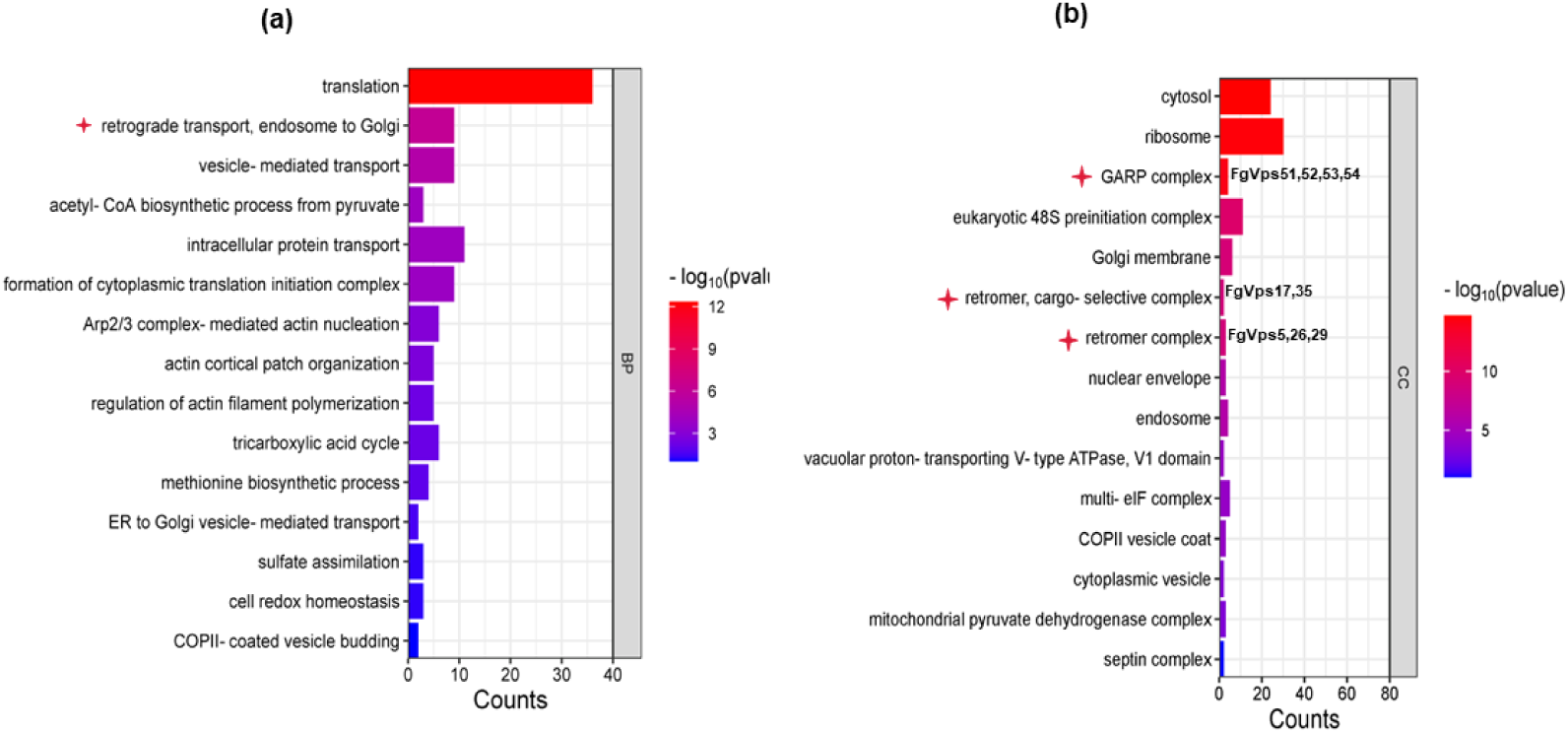
GO function analysis histogram. (a) The 15 most enriched GO pathways in Formula B (ranking by P value). BP: biological process. Asterisks represent the biological processes related to vesicle transport. (b) The 15 most enriched GO pathways in Formula B (ranking by P value). Cellular component (CC). Asterisks indicate the cellular component of retrograde transport, endosome to Golgi.

**Fig. S12.**
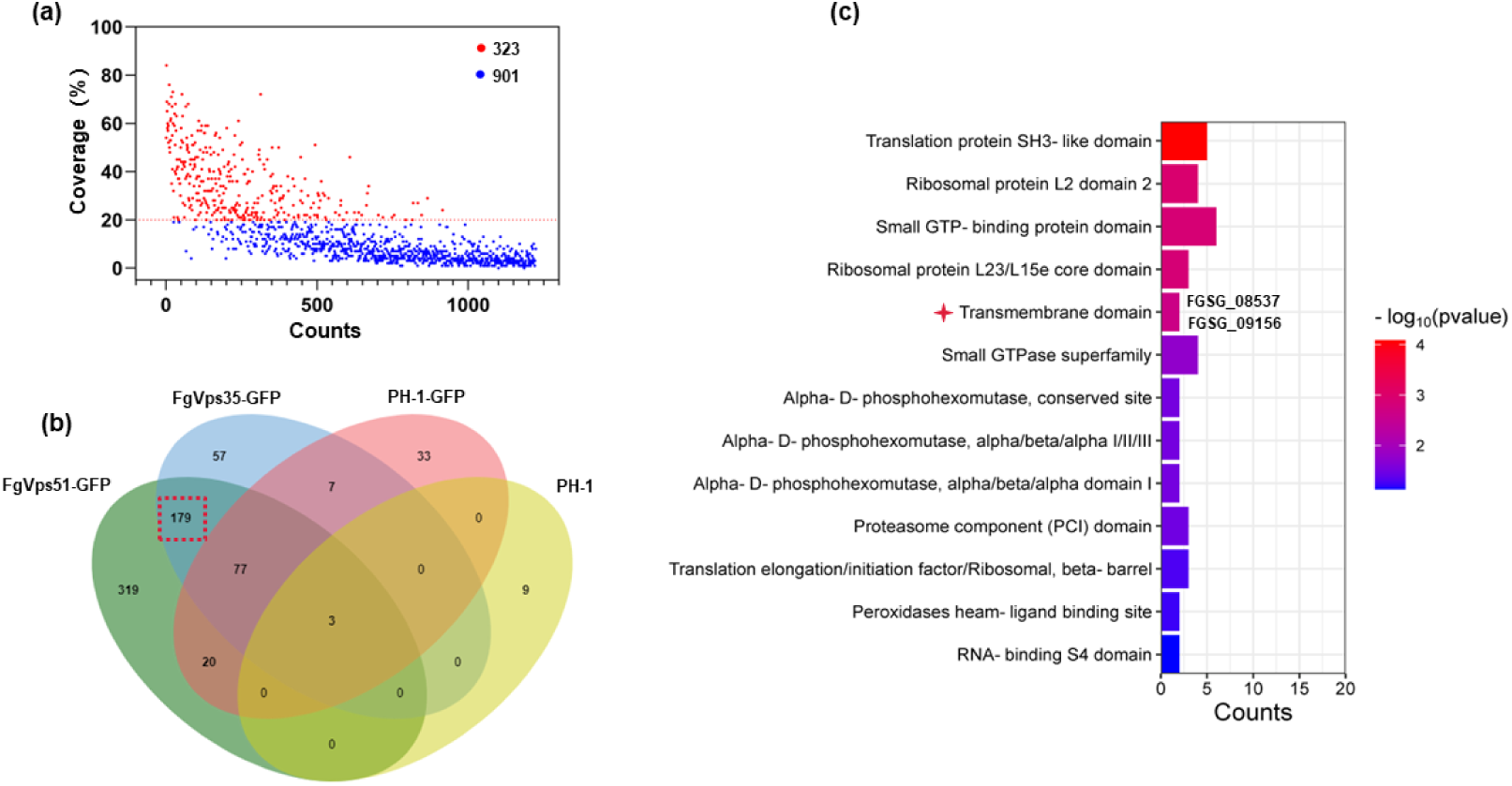
Identification of FgVps51- and FgVps35-interacting proteins. (a) 323 FgVps35-interacting partners were identified with 20% coverage and above. (b) In the Venn diagram of the FgVps35-GFP and FgVps51-GFP interaction groups, 197 proteins were specifically identified with high confidence, but not in the PH-1 and PH-1-GFP control groups. (c) Domain enrichment analysis.

**Fig. S13.**
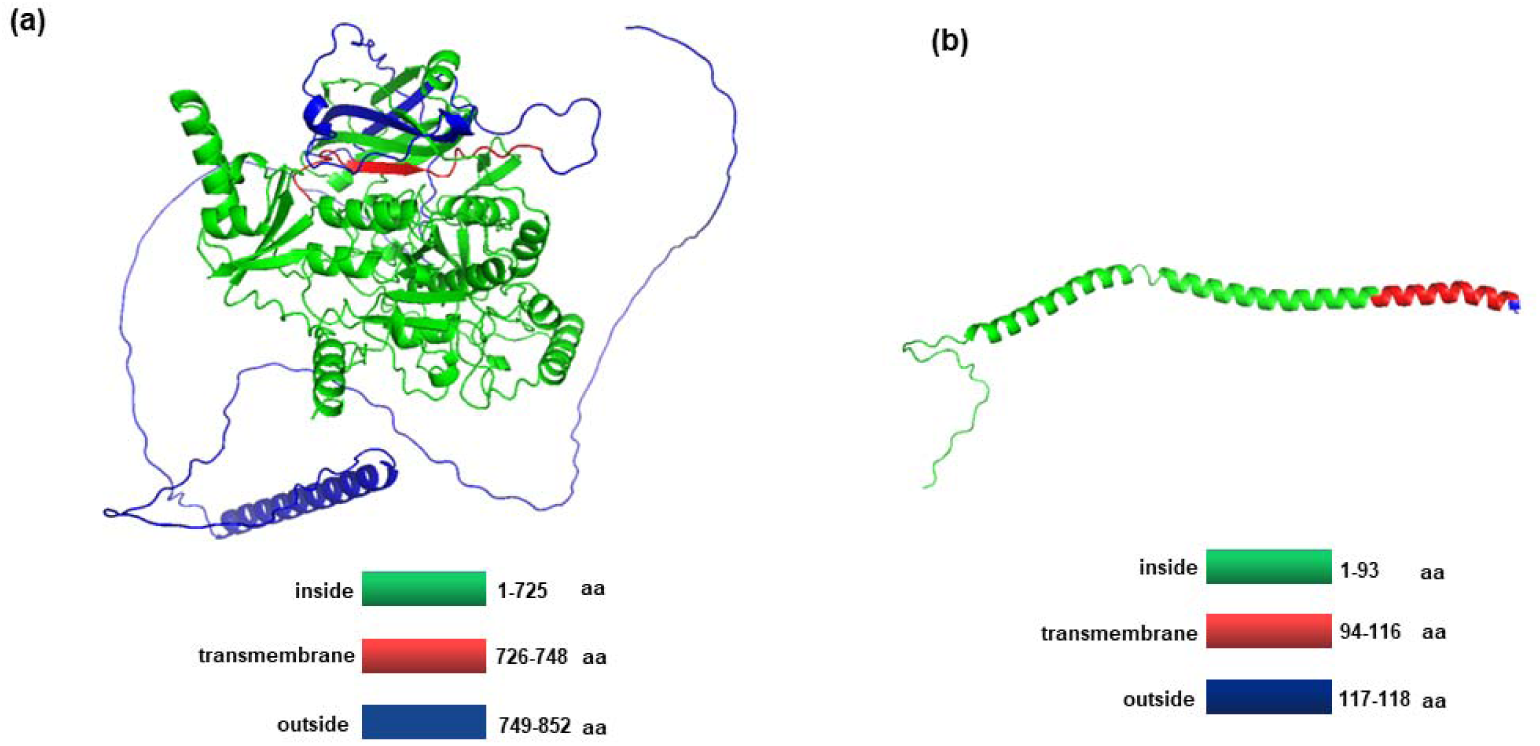
3D transmembrane domain structure of FgSnc1 and FgKex2 in *F. graminearum*. (a) The conserved TM (residues 94-116, red) domains of FgSnc1 (FGSG_08537) are shown and amino acid residues 1-93 are situated inside the membrane, while 100-112 residues are situated outside the membrane. (b) The conserved TM (residues 726-748, red) domains of FgKex2 (FGSG_09156) are shown and 1-725 amino acid residues are situated inside the membrane, while amino acid residues 749-852 are situated outside the membrane. The protein structures of FgSnc1 and FgKex2 were predicted with A_LPHA_F_OLD2_.

**Fig. S14.**
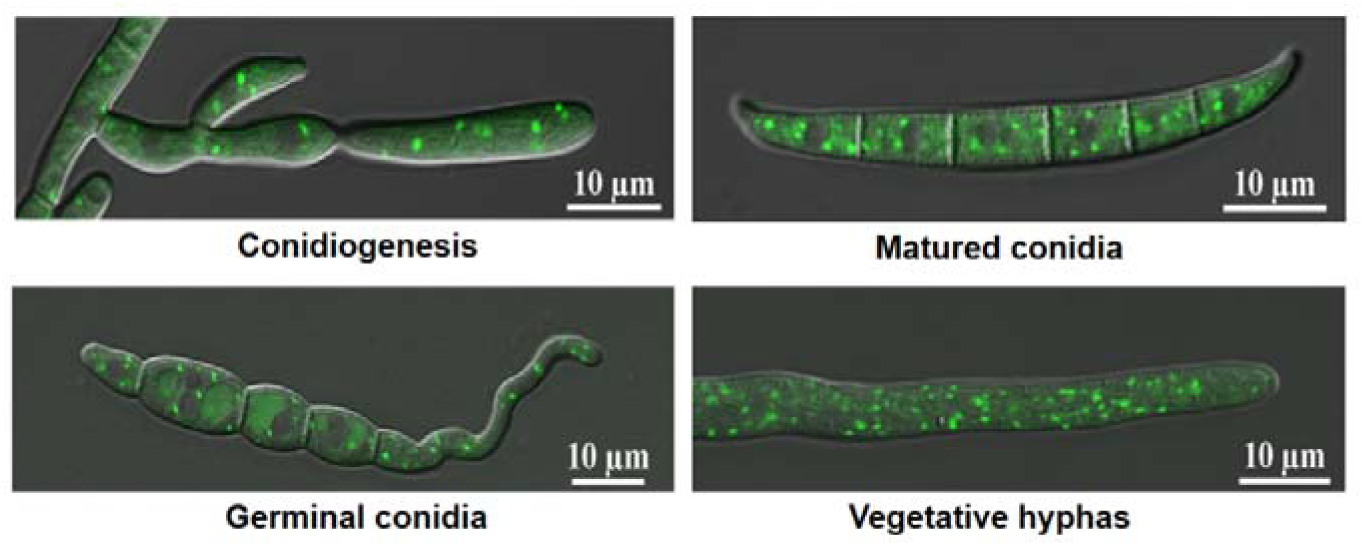
Subcellular localization of FgKex2-GFP. The expressions of FgKex2-GFP at different developmental stages were captured by Nikon TiE inverted laser scanning confocal microscope system. The fusion protein is localized in the cytoplasm of phialides cells and primary conidia, mature conidia, germinated conidia and vegetative hyphae as small dots.

**Fig. S15.**
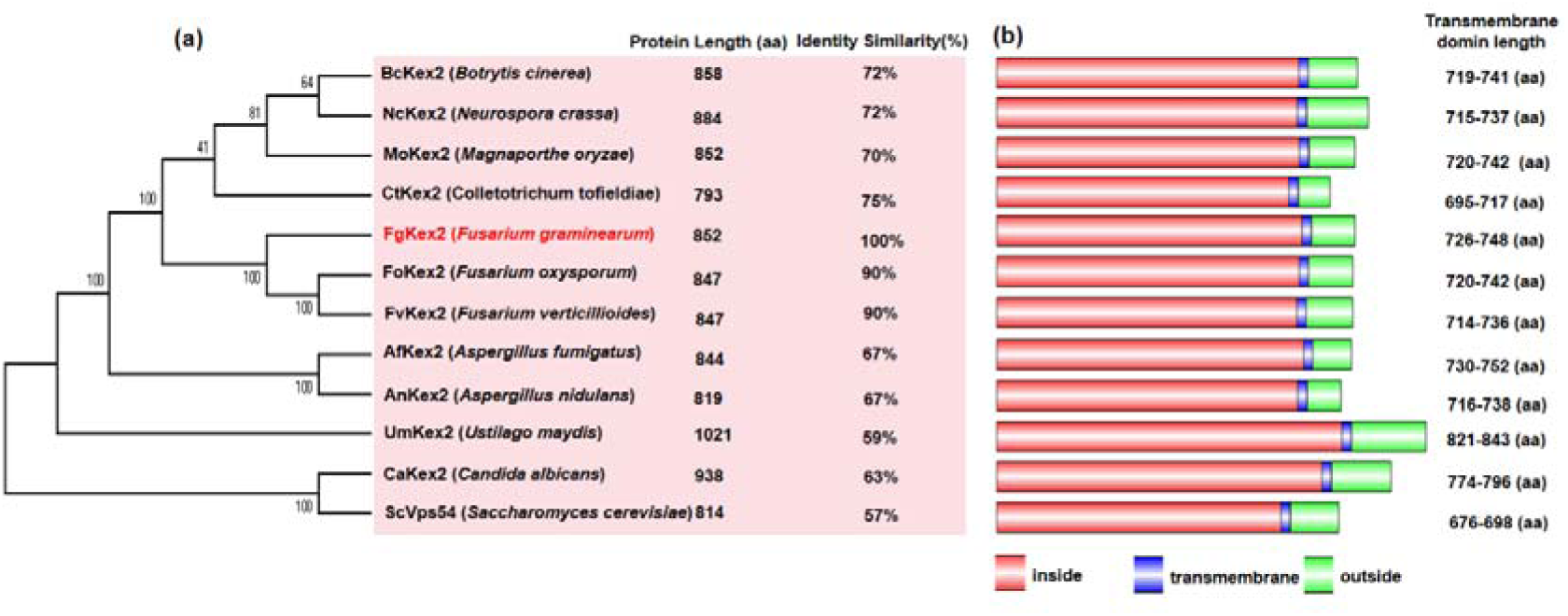
Phylogenetic and transmembrane domain structure analysis of Kex2 proteins from different species. (a) The analysis was performed using MEGA7 Program. The GenBank accession numbers (corresponding species names) of the Kex2 orthologs are as follows: XP_001559940.1 (BcKex2 *Botrytis cinerea*), XP_011393111.1 (NcKex2 *Neurospora crassa*), XP_003716137.1 (MoKex2 *Magnaporthe oryzae*), GKT91877.1 (CtKex2 *Colletotrichum tofieldiae*), XP_011328629.1 (FgKex2 *Fusarium graminearum*), KAG7415359.1 (FoKex2 *Fusarium oxysporum*), XP_018747745.1 (FvKex2 *Fusarium verticillioides*), XP_751534.1 (AfKex2 *Aspergillus fumigatus*), XP_661187.1 (AnKex2 *Aspergillus nidulans*), XP_011389281.1 (UmKex2 *Ustilago maydis*), O13359.2(CaKex2 *Candida albicans*), AJT30590.1 (ScVps54 *Saccharomyces cerevisiae*). (b) Schematic representation of the identified transmembrane domain-containing proteins in different species.

**Fig. S16.**
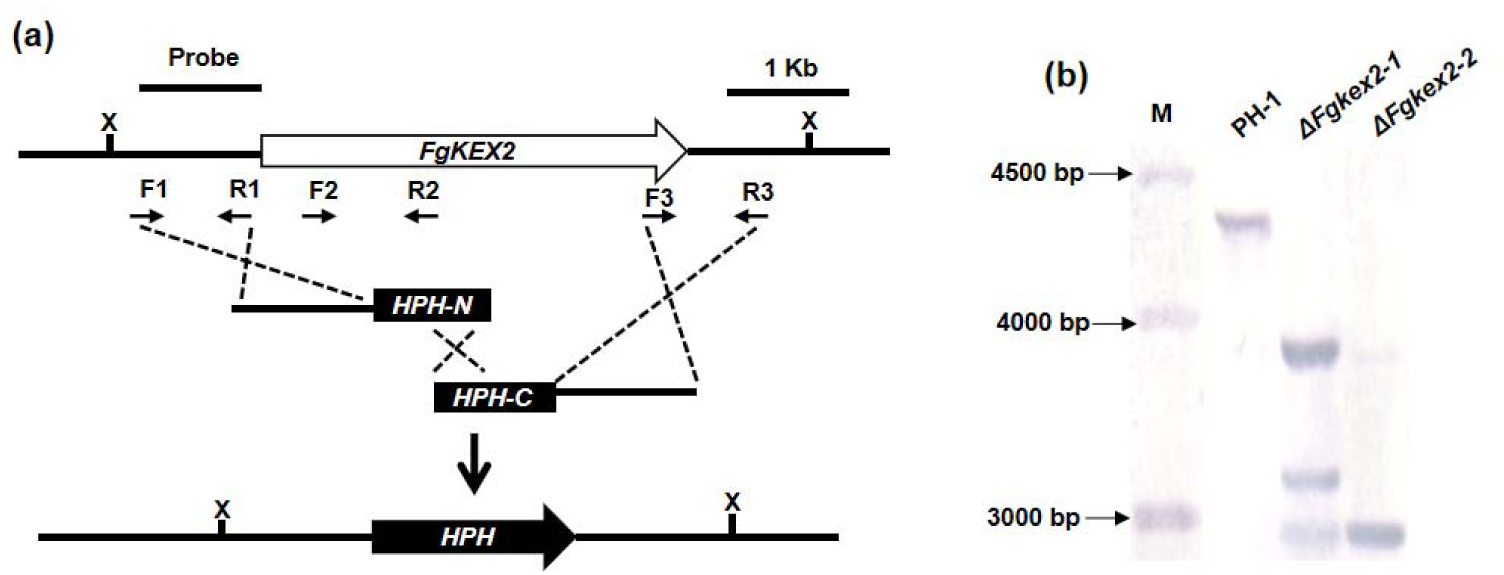
The targeted gene replacement strategy and Southern blot assay for *FgKEX2* gene deletion. (a) The *FgKEX2* targeted gene replacement strategy is shown in the schematic diagram. The primer pairs F1/R1 and F3/R3 were used to generate the gene replacement constructs. Primers F2 and R2 were used for mutant screening and identification. (b) Southern blot confirmation of *FgKEX2* gene deletion. *Xba*l-digested DNAs showed a 4.33 kb band in the WT and a 2.93 kb band in the mutants.

**Fig. S17.**
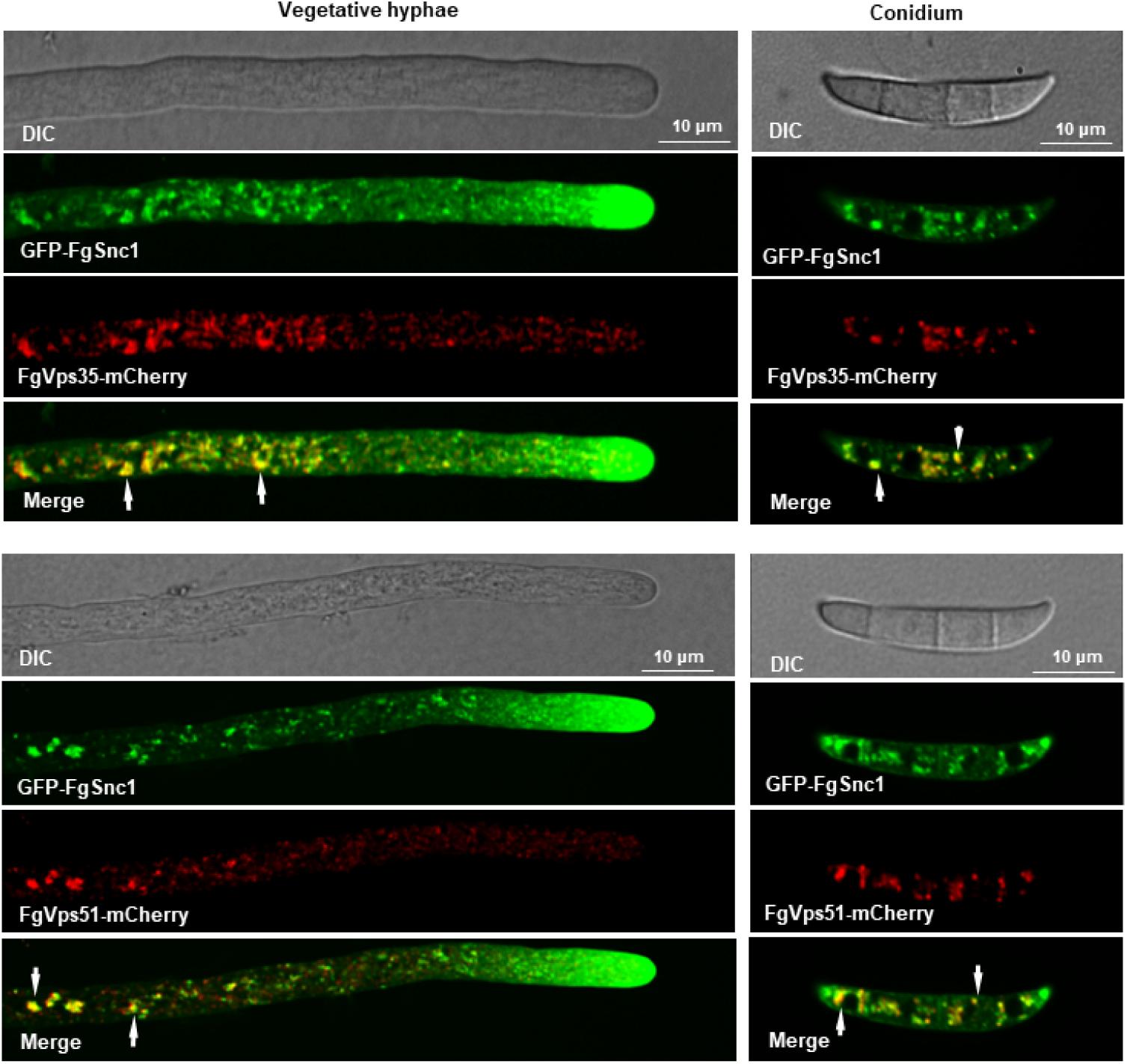
FgSnc1 interacts with GARP and retromer complexes. Representative confocal microscopy images showing partial co-localization of FgVps51-mCherry and FgVps35-mCherry with GFP-FgSnc1, respectively, in vegetative hyphae and conidia (yellow arrow).

**Fig. S18.**
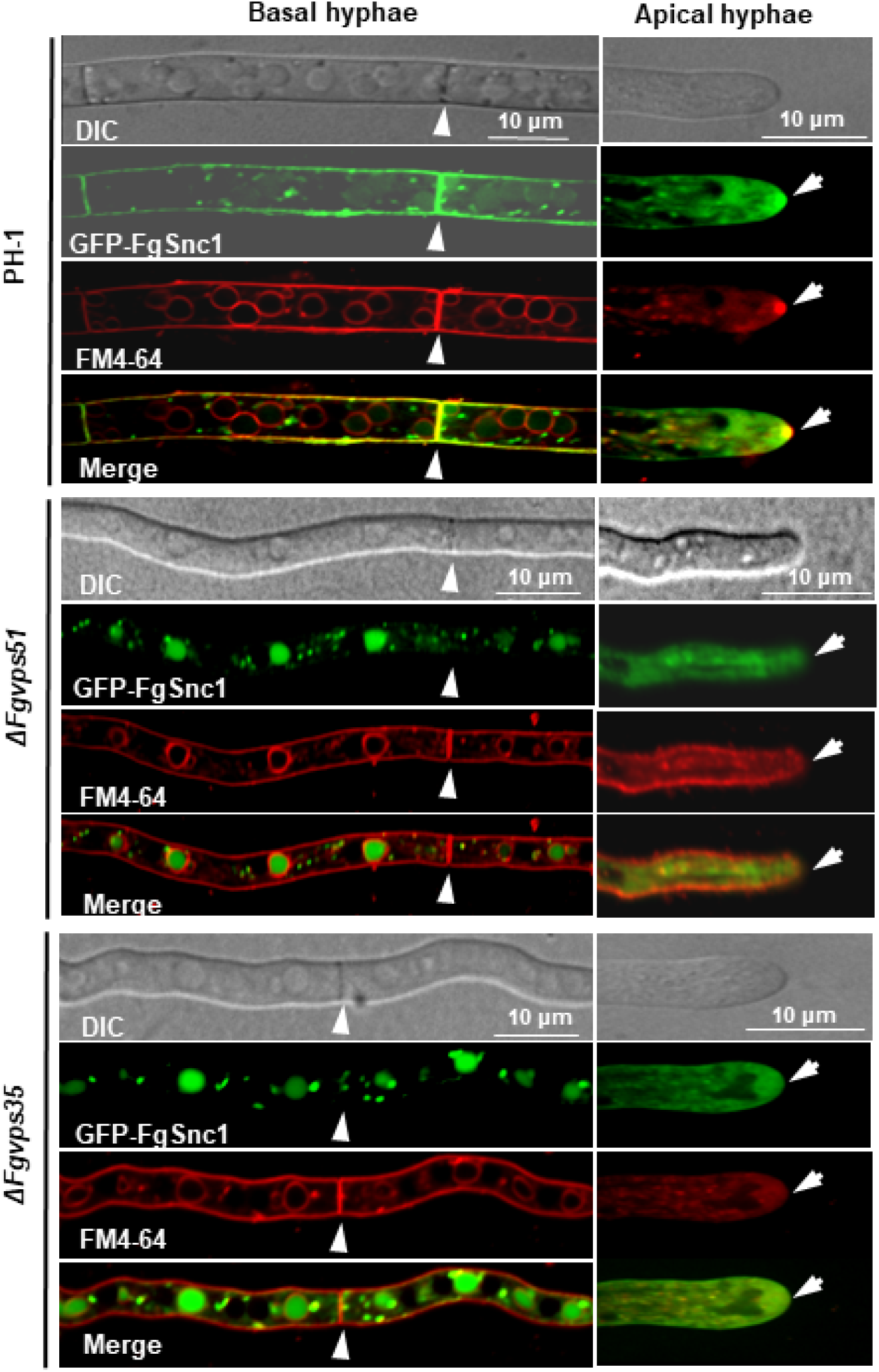
FgSnc1 is a cargo protein for the GARP/retromer-mediated vesicle trafficking pathway. Deletions of *FgVPS51* and *FgVPS35* perturb FgSnc1 transport to the plasma membrane and hyphal tip by mis-sorting FgSnc1 to the degradation compartments. In the WT hyphae, GFP-FgSnc1 was detected on the plasma membrane and hyphal tips. In contrast, GFP-FgSnc1 was mis-sorted into the vacuoles in the Δ*Fgvps51* and Δ*Fgvps35* mutants. Video S1 Co-motility of FgVps35-GFP and FgVps51-mCherry in hyphae of *F. graminearum*. Video S2 Co-motility of FgVps35-GFP and FgVps53-mCherry in hyphae of *F. graminearum*. Video S3 Co-motility of FgVps35-GFP and FgSec7-mCherry in conidia of *F. graminearum*. Video S4 Co-motility of FgKex2-GFP and FgSec7-mCherry in hyphae of *F. graminearum*. Video S5 Co-localization of GFP-FgSnc1 and FgSec7-mCherry in growing hyphae of *F. graminearum*.

**Table S1.**
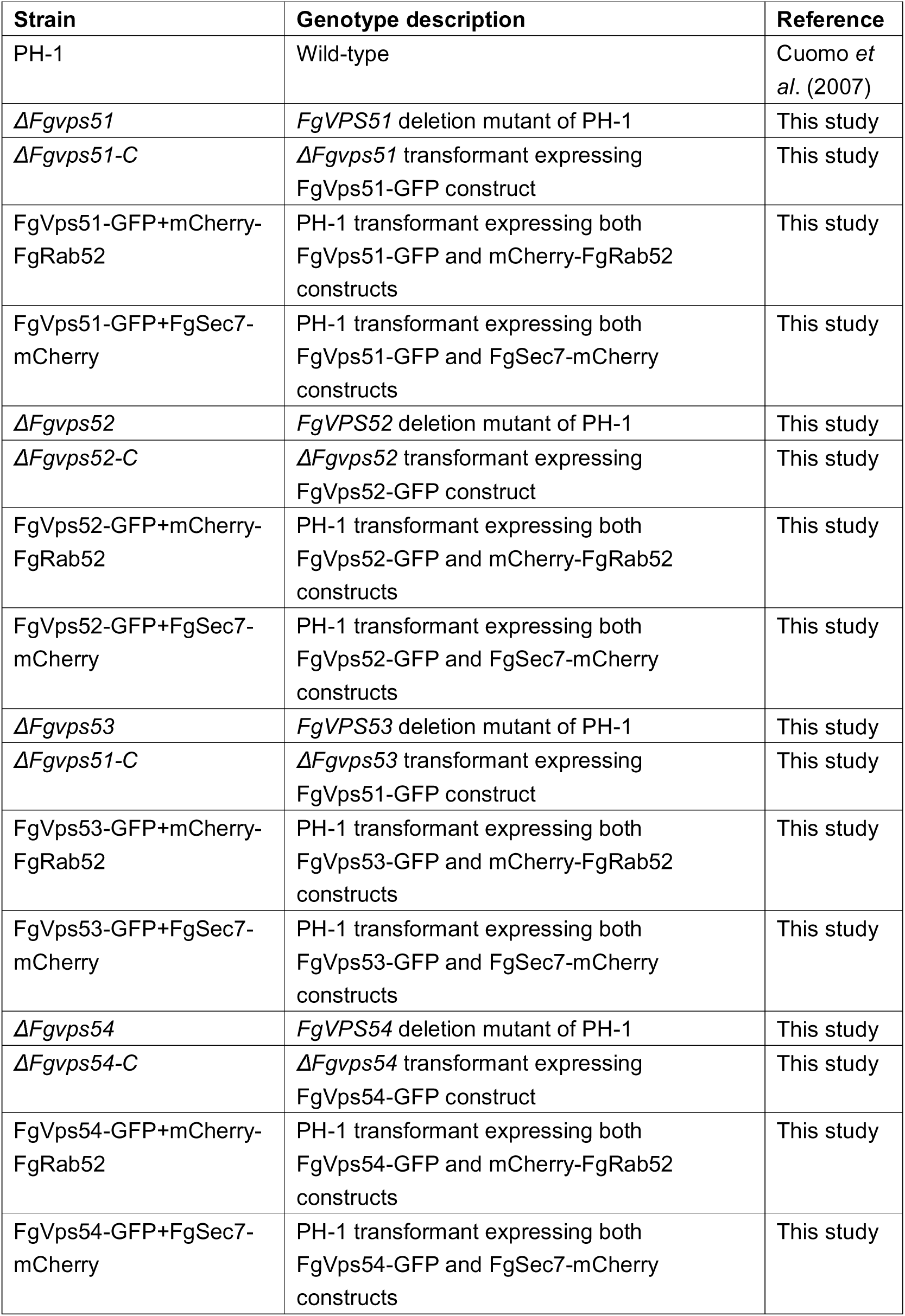
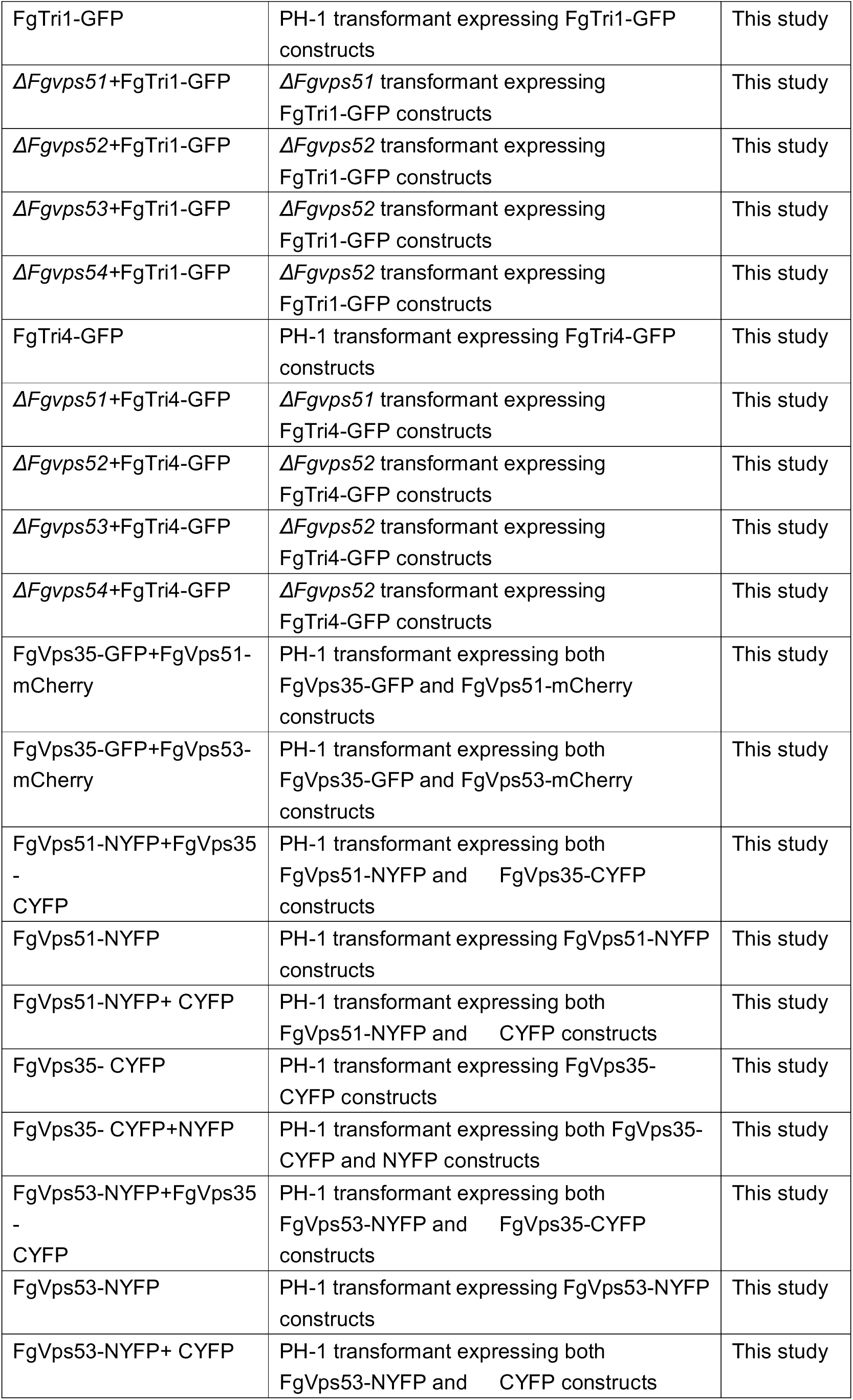
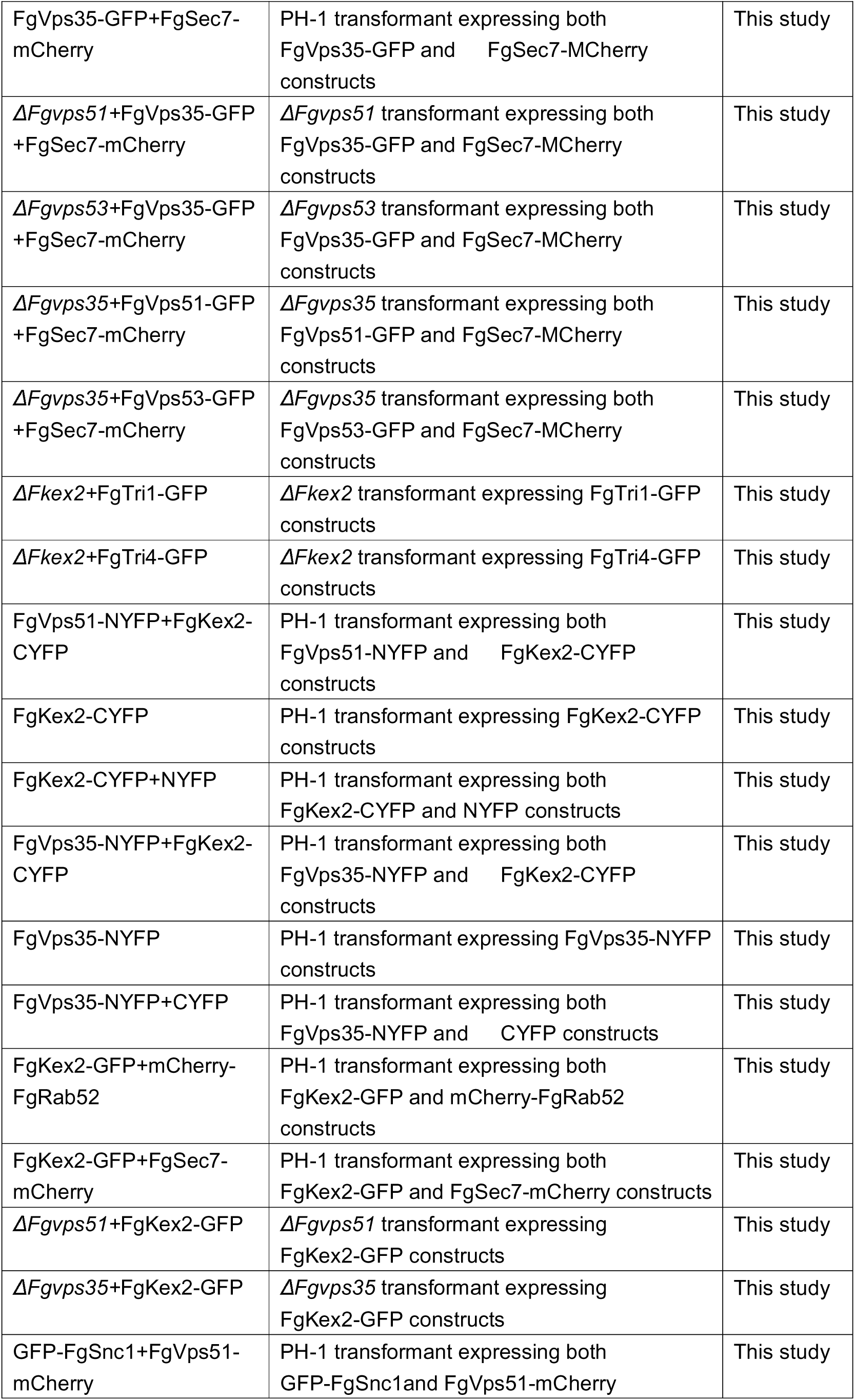
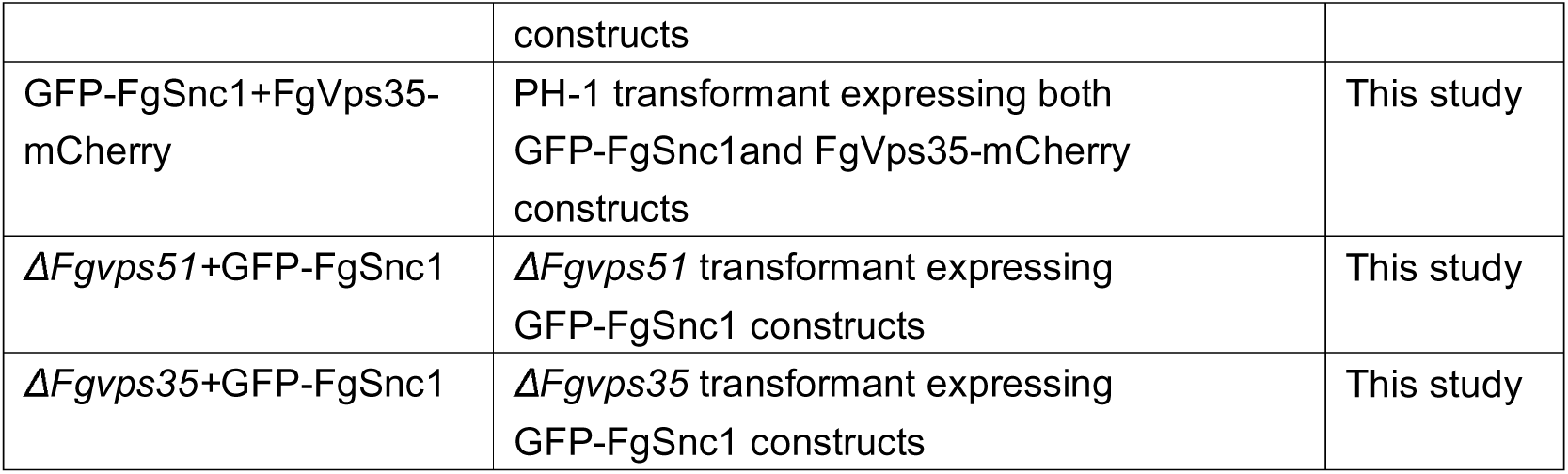
Wild-type and mutant strains of fungi used in this study.

**Table S2.**
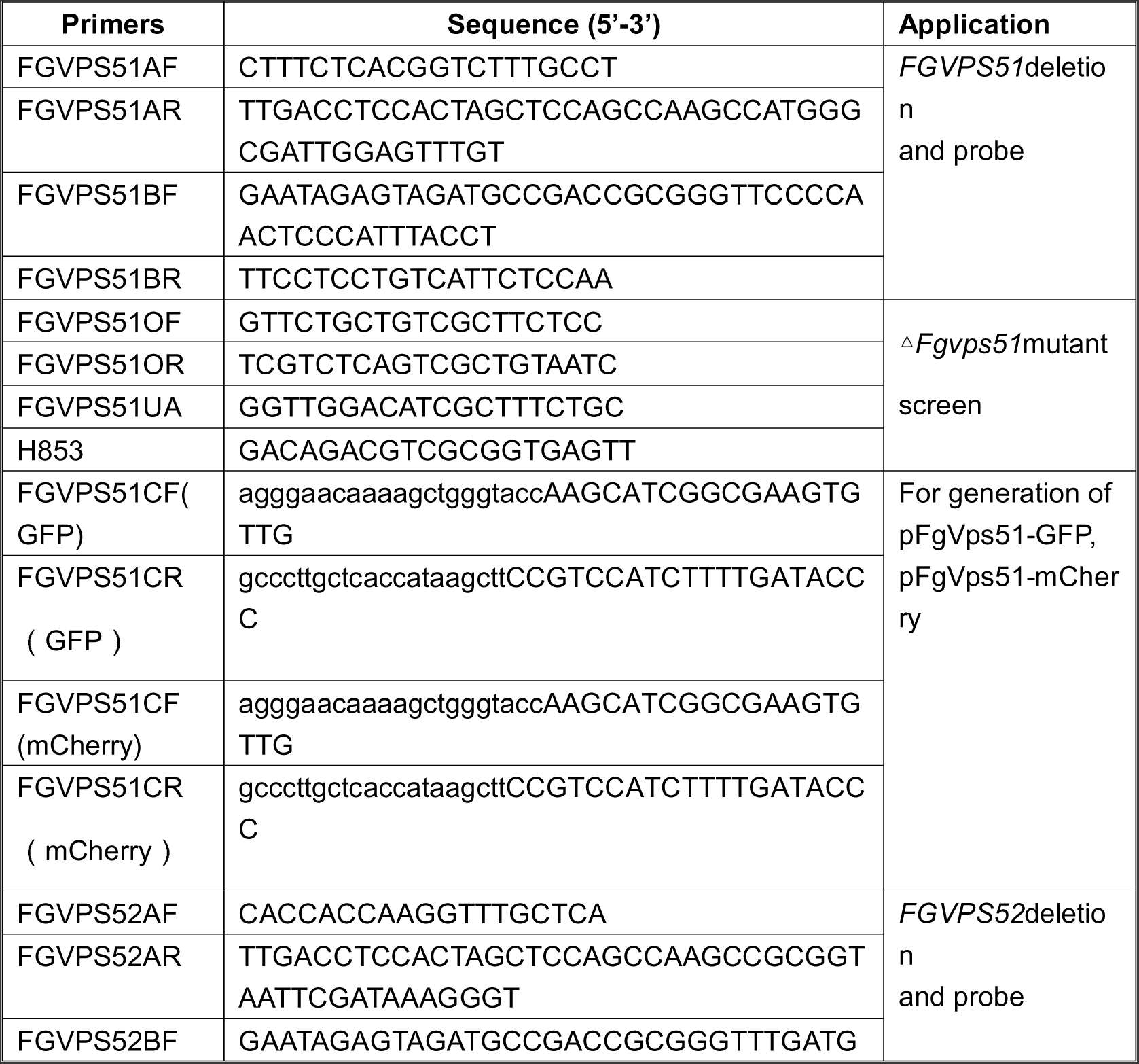
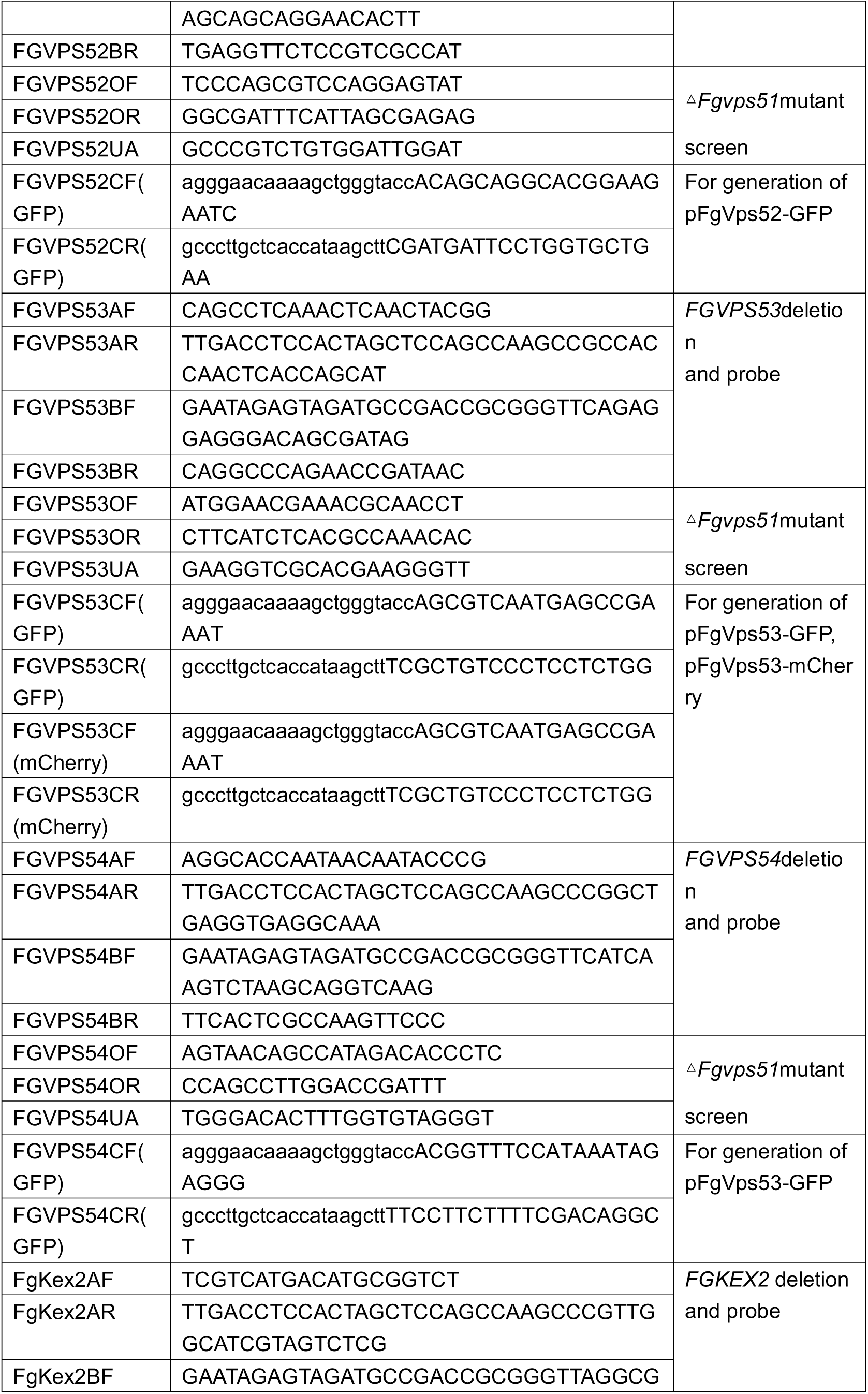
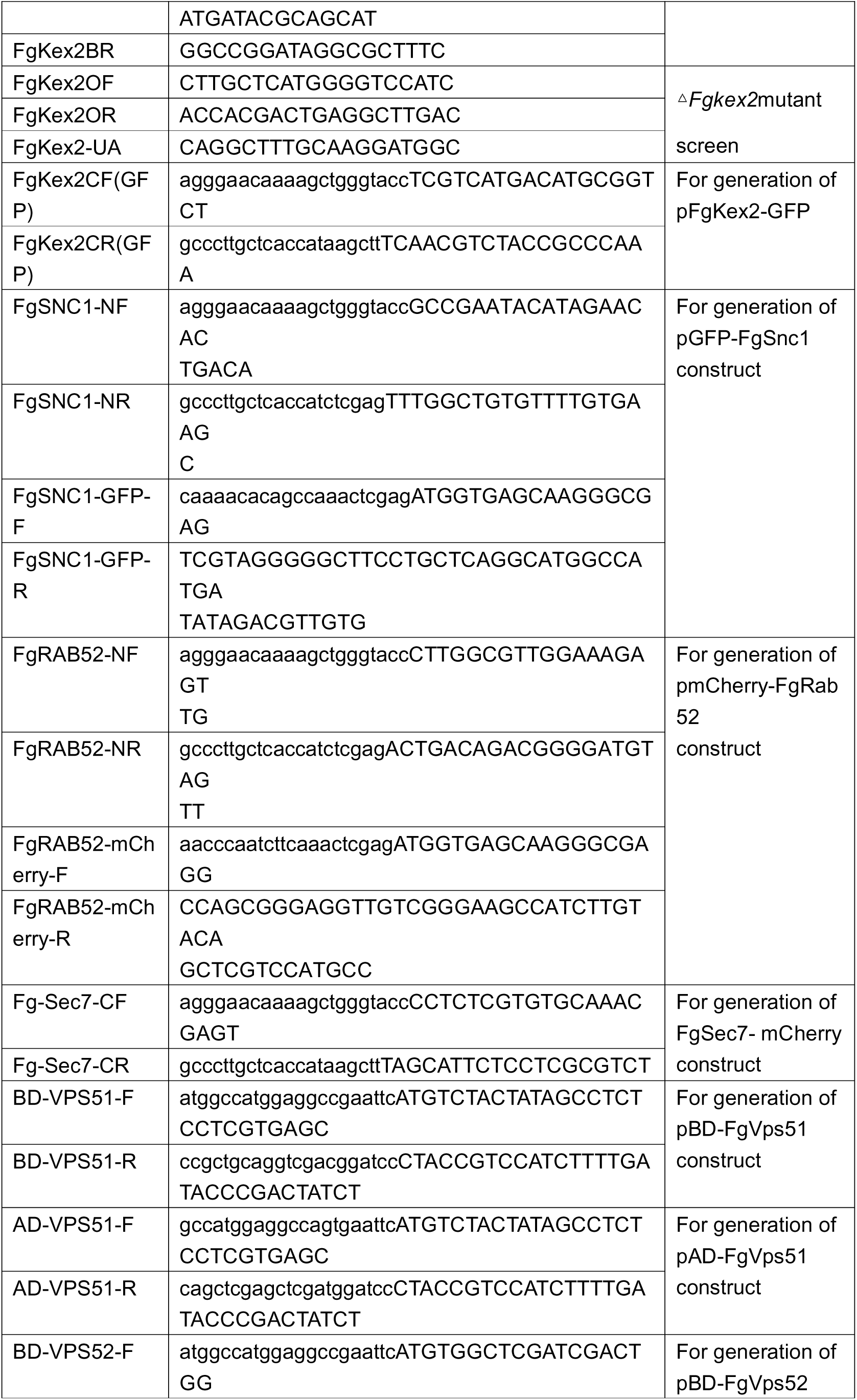
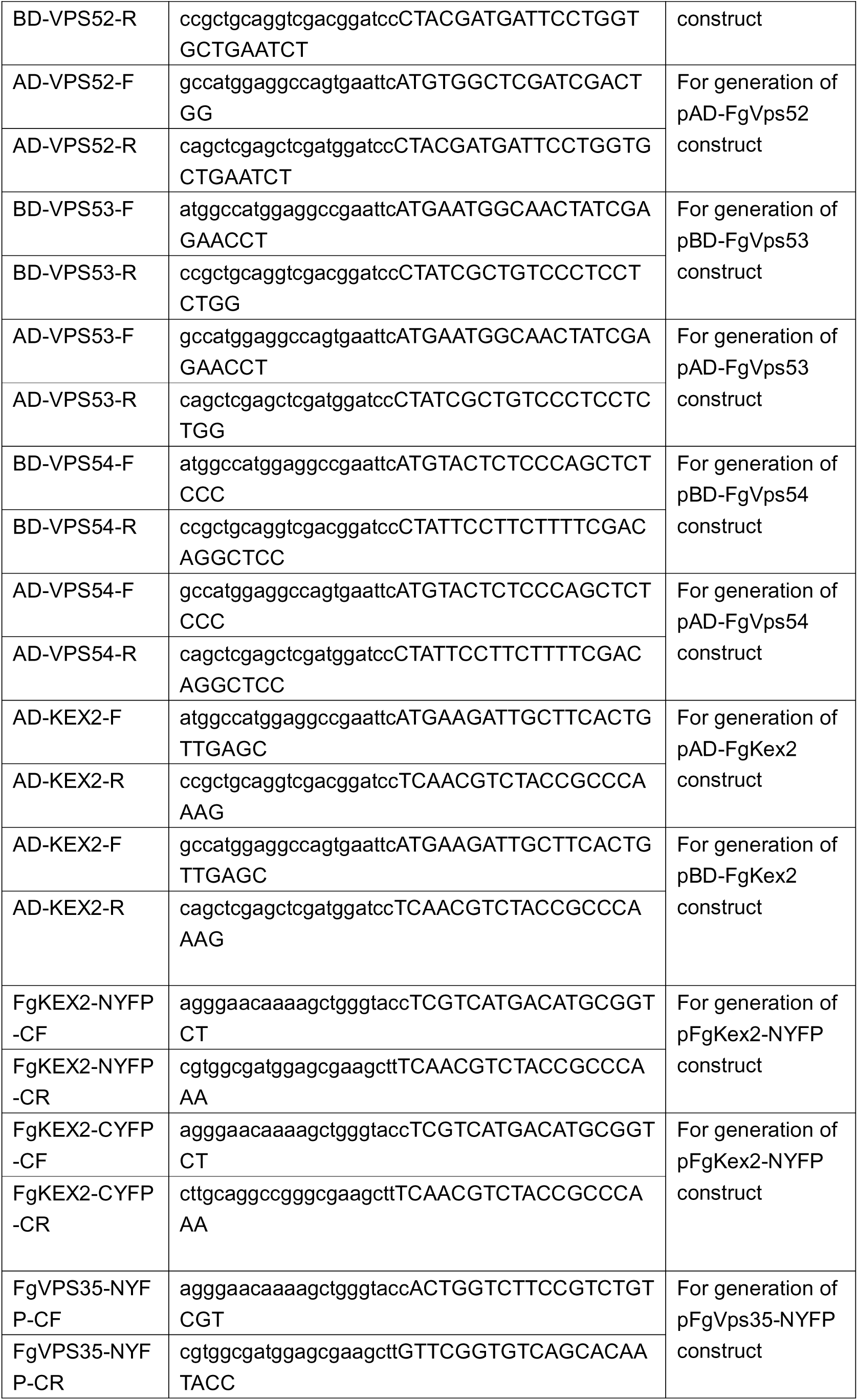
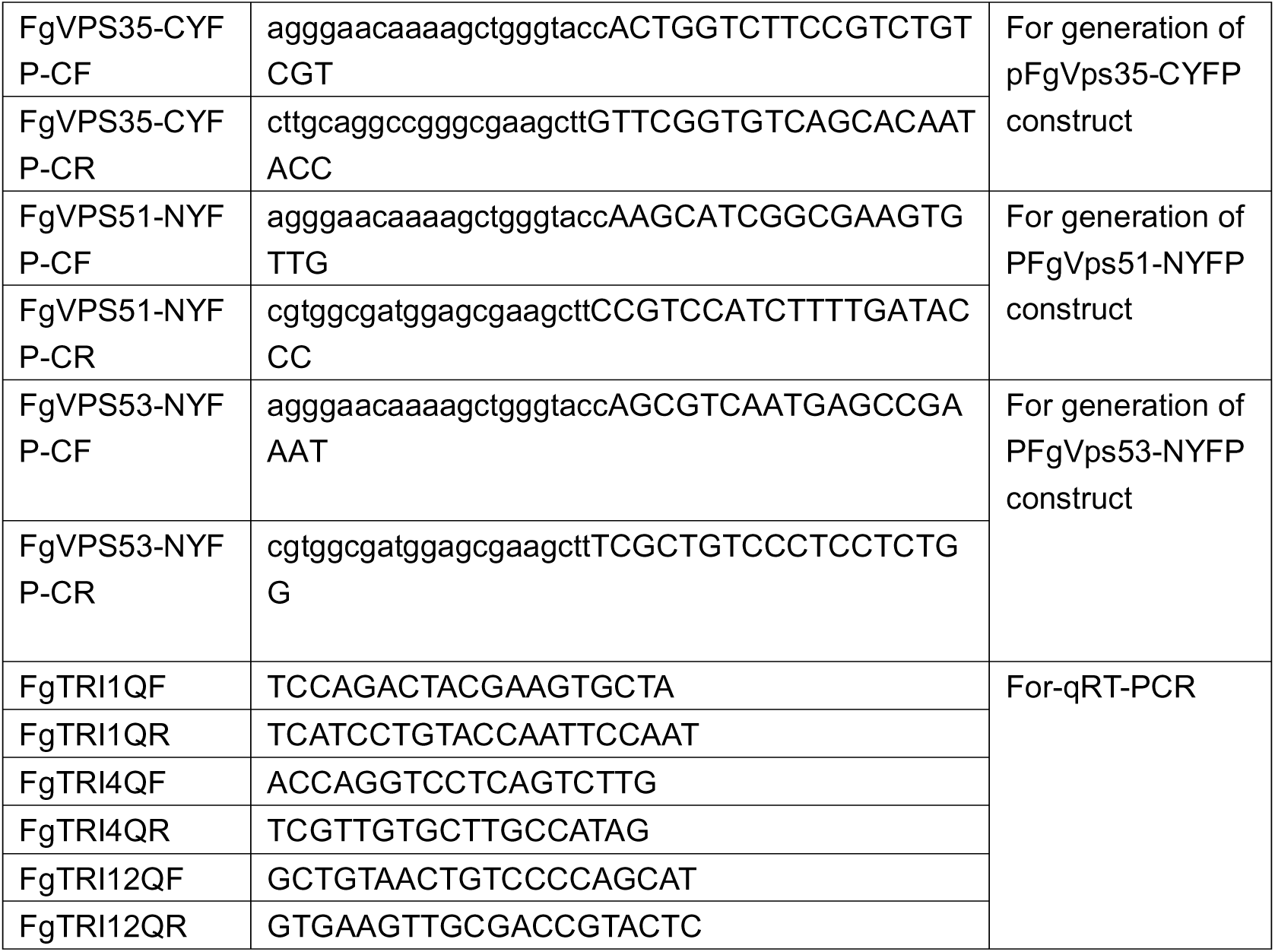
PCR primers used in this study.

**Table S3.** Plasmids used in this study.

| Clone | Description |
| --- | --- |
| pFgVps51-GFP | For expression of FgVps51: GFP, cloned in *KpnI-HindIII* sites of pKNT-GFP. Ampicillin and Neomycin resistance |
| pFgVps52-GFP | For expression of FgVps51: GFP, cloned in *KpnI-HindIII* sites of pKNT-GFP. Ampicillin and Neomycin resistance |
| pFgVps53-GFP | For expression of FgVps51: GFP, cloned in *KpnI-HindIII* sites of pKNT-GFP. Ampicillin and Neomycin resistance |
| pFgVps54-GFP | For expression of FgVps51: GFP, cloned in *KpnI-HindIII* sites of pKNT-GFP. Ampicillin and Neomycin resistance |
| pFgVps51-mCheery | For expression of FgVps51: mCheery, cloned in *KpnI-HindIII* sites of pKNT- mCheery. Ampicillin and Neomycin resistance. |
| pFgVps53-mCheery | For expression of FgVps53: mCheery, cloned in *KpnI-HindIII* sites of pKNT- mCheery. Ampicillin and Neomycin resistance. |
| pFgSec7-mCheery | For expression of FgSec7: mCheery, cloned in *KpnI-HindIII* sites of pKNT- mCheery. Ampicillin and Neomycin resistance. |
| pmCherry-FgRab52 | For expression of mCherry: FgRab52, cloned in *KpnI-BamHI* sites of pKNT. Ampicillin and Neomycin resistance |
| pFgKex2-GFP | For expression of FgKex2: GFP, cloned in *KpnI-HindIII* sites of pKNT-GFP. Ampicillin and Neomycin resistance |
| pFgVps35-GFP | For expression of FgVps35: GFP, cloned in *KpnI-HindIII* sites of pKNT-GFP. Ampicillin and Neomycin resistance |
| pFgSnc1-GFP | For expression of GFP: FgSnc1, cloned in *KpnI-BamHI* sites of pKNT. Ampicillin and Neomycin resistance |
| pFgVps51-NYFP | For expression of FgVps51: NYFP, cloned in *KpnI-HindIII* sites of pKNT-NYFP. Ampicillin and Neomycin resistance |
| pFgVps53-NYFP | For expression of FgVps53: NYFP, cloned in *KpnI-HindIII* sites of pKNT-NYFP. Ampicillin and Neomycin resistance |
| PFgVps35-CYFP | For expression of FgVps35: CYFP, cloned in *KpnI-HindIII* sites of pCX62-CYFP. Ampicillin and Hygromycin resistance. |
| PFgKex2-CYFP | For expression of FgKex2: CYFP, cloned in *KpnI-HindIII* sites of pCX62-CYFP. Ampicillin and Hygromycin resistance. |
| pBD-FgVps51 | FgVps51 was cloned in pDHB1. Kanamycin resistance. |
| pBD-FgVps52 | FgVps52 was cloned in pDHB1. Kanamycin resistance. |
| pBD-FgVps53 | FgVps53 was cloned in pDHB1. Kanamycin resistance. |
| pBD-FgVps54 | FgVps54 was cloned in pDHB1. Kanamycin resistance. |
| pBD-FgKex2 | FgKex2 was cloned in pDHB1. Kanamycin resistance. |
| pAD-FgVps51 | FgVps51 was cloned in pPR3-N. Ampicillin resistance |
| pAD-FgVps35 | Obtained from Zheng et. al., 2016 |
| pAD-FgVps29 | Obtained from Zheng et. al., 2016 |
| pAD-FgVps26 | Obtained from Zheng et. al., 2016 |
| pAD-FgVps17 | Obtained from Zheng et. al., 2016 |
| pAD-FgVps5 | Obtained from Zheng et. al., 2016 |

**Table S4.**
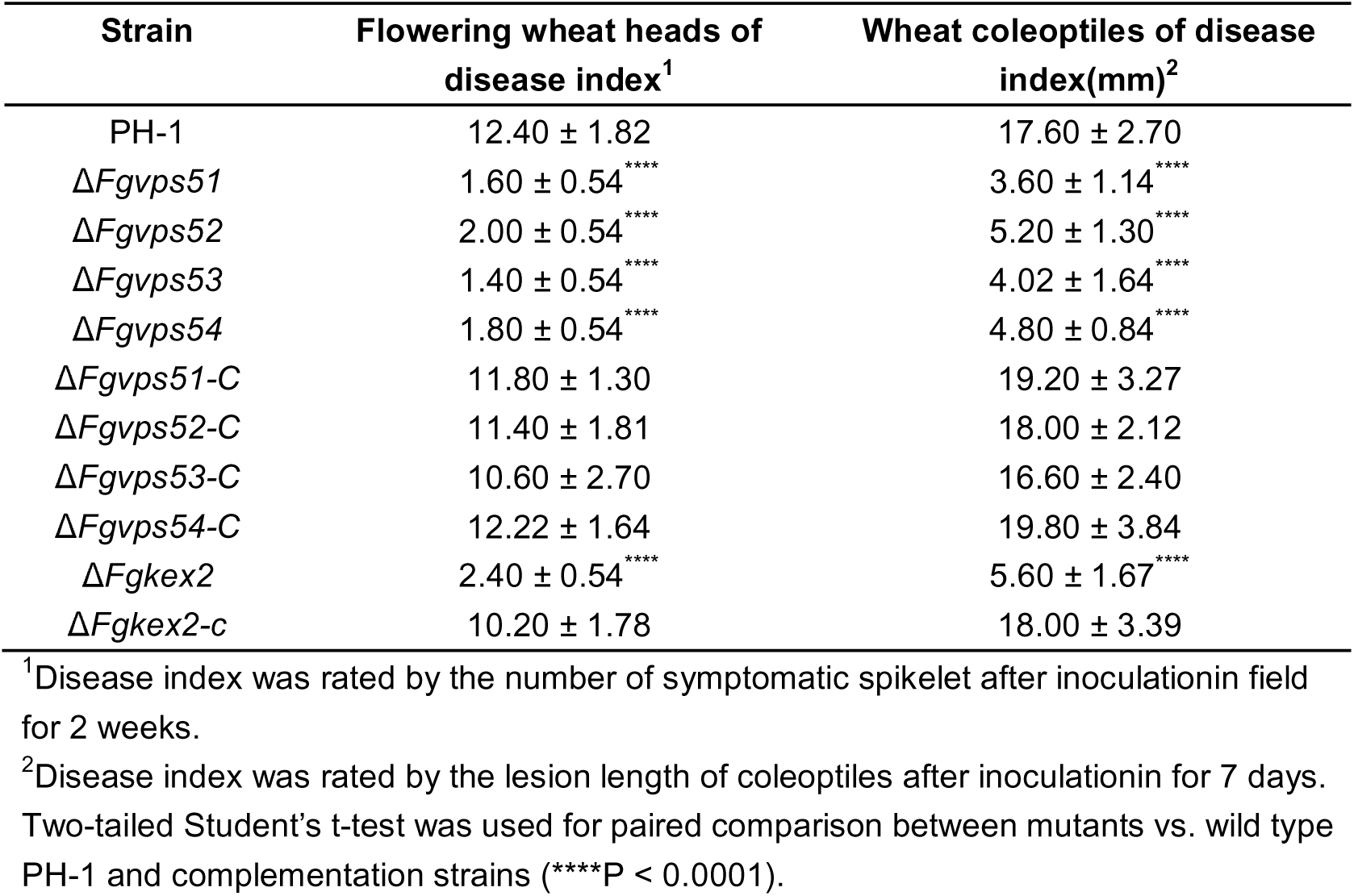
Characterization of disease inde.

